# Axial motion estimation and correction for simultaneous multi-plane two-photon calcium imaging

**DOI:** 10.1101/2021.09.28.462125

**Authors:** Andres Flores Valle, Johannes D. Seelig

## Abstract

Two-photon imaging in behaving animals is typically accompanied by brain motion. For functional imaging experiments, for example with genetically encoded calcium indicators, such brain motion induces changes in fluorescence intensity. These motion related intensity changes or motion artifacts cannot easily be separated from neural activity induced signals. While lateral motion within the focal plane can be corrected by computationally aligning images, axial motion, out of the focal plane, cannot easily be corrected.

Here, we develop an algorithm for axial motion correction for non-ratiometric calcium indicators taking advantage of simultaneous multi-plane imaging. Using at least two simultaneously recorded focal planes, the algorithm separates motion related and neural activity induced changes in fluorescence intensity. The developed motion correction approach allows axial motion estimation and correction at high frame rates for isolated structures in the imaging volume *in vivo,* such as sparse expression patterns in the fruit fly brain.

## Introduction

Using optical microscopy for the investigating of neural circuits *in vivo,* and particularly in behaving animals, is complicated by movement of the brain with respect to the focal plane^1–5^. In many preparations, such as mice or fruit flies, movement is reduced by mechanically stabilizing the brain, which however often still leaves residual motion^2^ and additionally can be invasive^6,7^.

In in vivo preparations, the brain moves in all three spatial dimensions, laterally, within the focal plane, as well as axially, out of the focal plane. Since the lateral resolution of the microscope is higher than the axial resolution, lateral motion is easier to detect^5^. Lateral motion can be corrected by post processing with a range of different methods, resulting in sequences of images aligned to a common field of view^1–3,5,8,9^.

Motion in axial direction, however, can usually not be corrected using post-processing. Axial motion leads to variations in the recorded fluorescence intensity due to changes in sample excitation. Generally, such changes cannot be disentangled from intensity fluctuations of the indicator resulting from neural activity^4^. Additionally, sample movement, which for example in mice lies in the range of a few micrometers^2,4^, can exceed the focal range of the two-photon microscope^10^ and therefore lead to loss of information^11^.

Algorithms similar to those used for lateral motion correction have also been applied for time series of volume images (z-stacks). Recording time series of *z*-stacks with sufficient axial resolution however limits the frame rate in each focal plane^12,13^. Faster axial correction speeds can be achieved by actively tracking moving brain samples. Taking advantage of a reference signal, for example a reflected beam^14^, behavioral parameters^15^, or fluorescent fiducials such as cell bodies^4,16^ or fluorescent beads^4^, the focal plane or additionally the region of interest (ROI) can be updated in a closed-loop configuration. Repositioning of the ROI at high frame rates and in fast control cycles however requires specialized hardware allowing random access scanning^4^. Additionally, introducing fiducials, typically fluorescent beads, is often impractical and invasive, for example when imaging in the brain of *Drosophila*.

Similarly, naturally occurring fiducials such as cell bodies are often not present in the imaging volume. Further, often only a single fluorescent dye, for instance a non-ratiometric calcium indicator, is used.

Alternatively to recording volume images in *z*-stacks, simultaneous multi-plane imaging allows recording a limited number of focal planes at the same time^17–22^. Similar to imaging in a single focal plane, multi-plane recordings are still subject to axial motion artifacts. However, the correlated intensity changes across simultaneously recorded focal planes that result from sample motion, allow inferring motion information. This has been used for tracking fluorescent beads using temporally mutliplexed and axially offset beams^23–25^.

Here, we develop an approach for axial motion estimation and correction in the brain for two-photon functional imaging with non-ratiometric calcium indicators. The approach does not require fiducials and can be applied for isolated neural structures inside the imaging volume. The technique relies on an optimization algorithm that estimates axial brain motion as well as disentangles motion and neural activity related fluorescence changes based on simultaneous imaging in at least two focal planes. The algorithm is demonstrated using simulations with two and four simultaneously recorded focal planes, as well as validated using *in vivo* imaging in behaving fruit flies.

## Results

### Outline of motion correction approach

In simultaneous multi-plane imaging, motion along the axial direction displaces the sample at the same time in all focal planes. This correspondingly leads to related intensity changes in the different planes. For functional imaging with calcium indicators, changes in intensity due to sample motion are additionally confounded with neural activity related intensity changes. For disentangling these two contributions we developed an optimization based correction approach.

The experimental setup is shown in Fig. 1B and was previously described^22^. Simultaneous two-plane imaging was implemented with two temporally offset beams (by 6 ns) recorded independently using temporal multiplexing^17^. The axial profiles of the two beams are shown in Fig. 1C.

**Figure 1.**
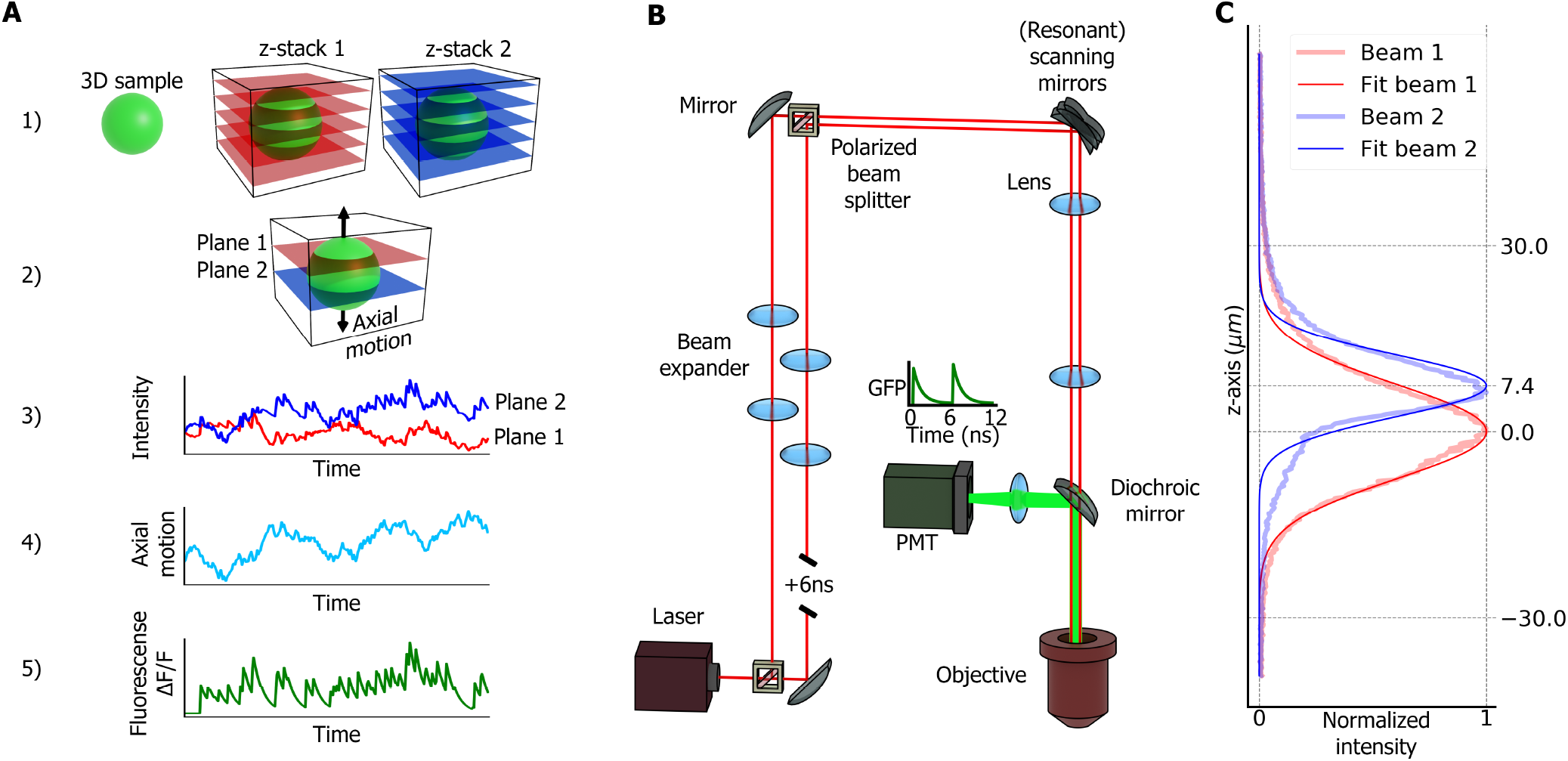
Approach and setup for motion estimation and correction. **A** Outline of method: 1) Calibration step: two stacks of the sample are recorded simultaneously in two different, axially offset focal planes. 2) Fluorescence intensity of the sample is recorded simultaneously in two planes and the sample moves in axial direction (*z*-axis). 3) Changes in intensities over time in each plane have two contributions: axial motion as well as neural activity. 4) The algorithm uses the recorded stacks to estimate axial motion of the sample from intensities recorded in the two planes. 5) Changes in fluorescence, Δ*F/F*, are corrected to remove the contribution of axial motion, yielding motion corrected, neural activity related fluorescence changes. **B** Optical setup: two Gaussian beams are temporally offset by 6 ns, allowing simultaneous imaging in two different planes using temporal multiplexing. **C** Normalized profiles of the two beams along the *z*-axis fitted with Gaussian functions.

The different steps of the correction approach are outlined in Fig. 1A. First, a calibration stack of the sample is recorded with the two temporally multiplexed beams (Fig. 1A,1)). This stack is averaged over 10 recordings (see Methods for details) so that intensity variations due to motion are averaged out. For subsequent combined functional imaging and motion estimation, the sample is then approximately centered between the two beams such that it is always visible in both focal planes (Fig. 1A, 2)). After recording a time series (Fig. 1A, 3)) and computationally removing lateral motion, the correction algorithm extracts axial sample motion. Using the estimated axial motion and the *z*-stacks, the intensity recorded by each beam at each focal plane is finally corrected for motion (Fig. 1A, 5)).

The algorithm does not require a second activity-independent label for motion estimation, such as a fluorescent bead of a different color. It however relies on the assumption that neural activity induced fluorescence changes occur with a constant gain factor from baseline in axial direction (see Supplementary Fig. S1 for an illustrative example). Additionally, the method requires that the structure of interest is always visible in both focal planes and isolated, meaning that no other structures move in and out of the imaging volume. Depending on the sample and its typical motion, the separation between focal planes as well as the axial width of the beams therefore needs to be adjusted.

The algorithm finds and corrects axial motion by optimizing a cost function. The cost function is derived and explained in detail in Methods in several steps. First an analytical approximation for a one dimensional sample is developed to illustrate the workings of the algorithm before introducing the general case with multiple ROIs. Both, the one dimensional as well as the full algorithm are demonstrated in simulations. We then test the algorithm experimentally. First, we estimate controlled focal plane motion in a sample labeled with green fluorescent protein. Then, in addition to motion, we also estimate controlled variations in fluorescence intensity. Next, we correct neural activity recorded with a genetically expressed calcium indicator for controlled motion. Finally, the algorithm is applied for measuring and correcting actual brain motion.

### Motion correction in simulated data for a single voxel

The motion correction algorithm was first verified using simulated data (Fig. 2). For this purpose, we defined an arbitrary one dimensional sample extended along the optical axis (*z*-axis, Fig. 2A, top). We additionally defined two Gaussian beams with different beam profiles and intensities, similar to the ones used in the experiment (Fig. 2A, top). The profiles of the beams along the *z*-axis must cover the axial range of sample motion, keeping the sample always visible with both beams. At the beginning of the simulated experiment, a *z*-stack is computed for each beam (Fig. 2A, bottom, see Methods for details). The sample is assumed to move along the *z*-axis as well as change its intensity due to neural activity (Fig. 2B). We assume that activity changes occur homogeneously along the optical axis, that is, with a multiplicative factor with respect to the fluorescence baseline (see Supplementary Fig. S1).

**Figure 2.**
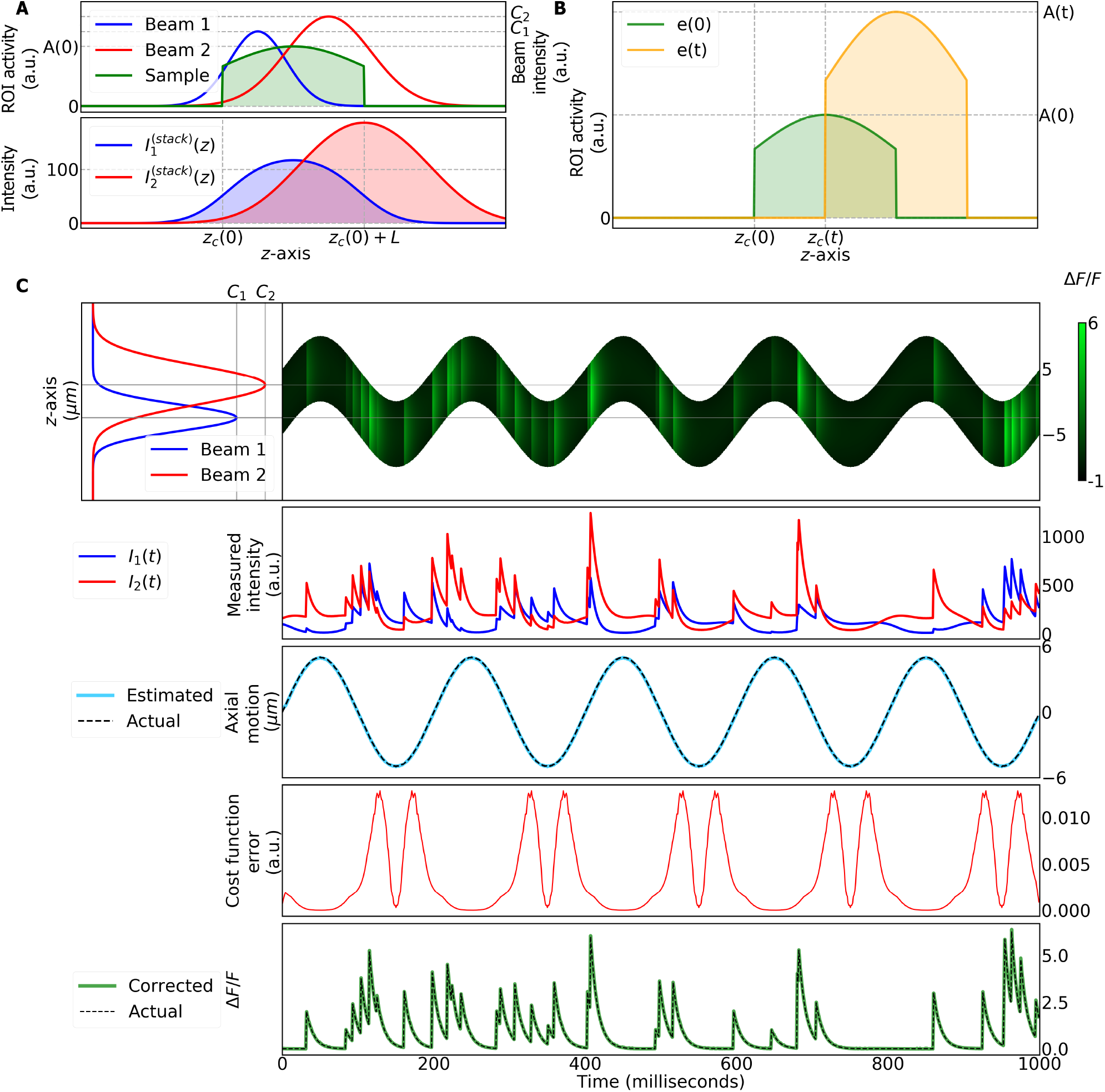
Motion correction in a simulated single ROI. **A** Top: axial intensity profiles of the two simulated beams and the simulated sample (ROI) at time *t* = 0. Bottom: two stacks recorded from the sample at time *t* = 0 with the two beams. **B** Example of axial motion and activity of the ROI. Sample profile along the *z*-axis at time *t* = 0 is shown in green. At time *t* > 0, the offset of the ROI along the *z*-axis changes while its activity increases. **C** Simulation of ROI activity and motion along the *z*-axis over time. Top row, left side: beam profiles. Right side: sample moves along *z*-axis over time. Color indicates sample neural activity. Second row: resulting intensities measured by each beam have two different contributions, motion and activity. Third row: comparison of actual and estimated axial motion. Fourth row: cost function error of the motion estimation algorithm (see Methods for details). Bottom row: actual and estimated, corrected changes in neural activity induced Δ*F/F*.

A simulated time series of sinusoidal axial sample motion combined with stochastic changes in neural activity is shown in Fig. 2C, top row. The resulting intensity time series, recorded simultaneously in two focal planes, are shown in Fig. 2C, second row. Based on the recorded intensity time series, the optimization algorithm accurately estimates both, sample motion as well as motion-corrected, activity related changes in fluorescence intensity (Fig. 2C, third and bottom rows).

### Motion correction in simulated data for multiple voxels

The algorithm was further validated with simulated data with multiple one dimensional voxels, representing regions of interests (ROIs) defined on a three dimensional sample (see next section and Methods for details). For this, neural activity in a ring attractor was simulated, similar to experimentally observed data^22,26^. A random distribution of the axial position of the different ROIs was assumed, reflecting structural variability (Fig. 3A, top row). Neural activity in this kind of network is centered in a bump and we here assumed that the bump of activity moves at constant velocity around the ring attractor. To simulate brain motion, the sample was oscillated with respect to the two focal planes along the axial direction (Fig. 3A, bottom). As for the case of a single voxel, a z-stack is computed using both beams. Intensity measurements from both beams are subjected to noise, similar to experimental recordings (see Methods for details). The algorithm assumes that all ROIs undergo the same axial motion. Therefore, all ROIs contribute to estimating motion (Fig. 3C, first row).

**Figure 3.**
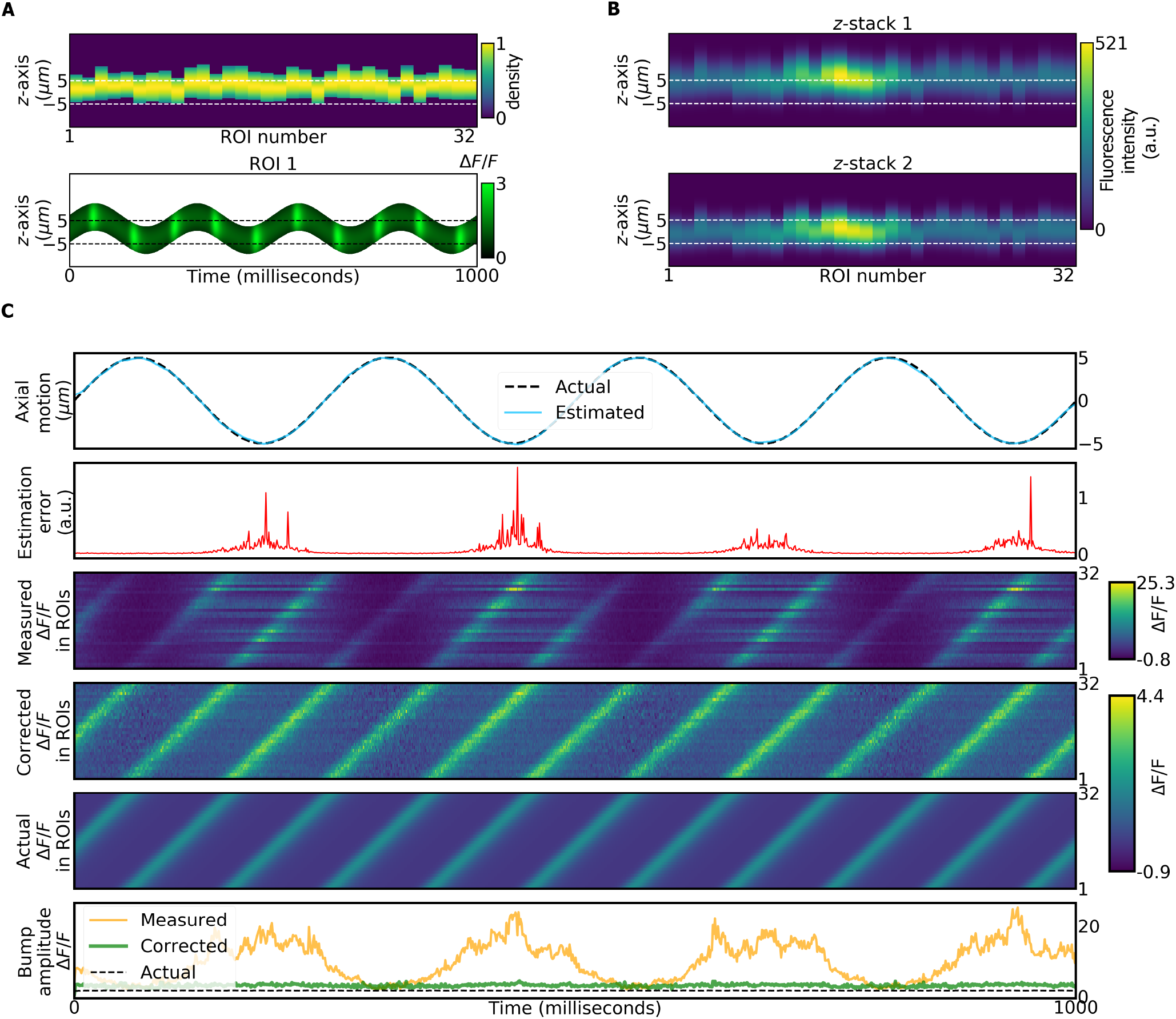
Motion correction in a simulated ring attractor with 32 ROIs. **A** Top: normalized intensity profiles of the ROIs, which are simulated with varying lengths and offsets along the *z*-axis. **B** Bottom: ROI 1 moves along the *z*-axis over time while its activity (ΔF/F) changes (see C for all ROIs). **B** Top: stacks of the ROIs obtained with first beam. Bottom: same for second beam. **C** Simulation of ROIs with axial motion and activity changes over time. All ROIs move together, while activity changes independently in each ROI. First row: comparison of actual and estimated axial motion of ROIs. Second row: cost function error of the motion estimation algorithm. Third row: measured changes in fluorescence with combined activity changes and axial motion. Fourth row: changes in fluorescence after motion correction. Fifth row: actual changes in fluorescence due to activity. Bottom row: comparison of the averaged measured, corrected and actual bump amplitudes in the ROIs (see Methods for details on all steps).

The bump of activity traveling with continuous velocity across the different ROIs is shown before correction in Fig. 3C, third row, as well as after correction in Fig. 3C, fourth row, and compared with the actual bump activity in Fig. 3C, fifth row. As also seen in the average over all ROIs (Fig. 3C, bottom row), the algorithm accurately corrects motion artifacts in the estimation of fluorescence changes.

### Estimation and correction of controlled sample motion *in vivo*

The motion correction algorithm was experimentally tested by controlled variation of the focal plane in neurons labeled with green fluorescent protein (GFP, Figs. 4 and 5). For this, GFP was expressed in wedge neurons of the head direction system of the fruit fly^26^ (R60D05-GAL4, Fig. 4A, left side). The fly’s proboscis was fixed with wax to prevent brain motion^7^. To simulate sample motion, the axial position of the microscope objective was modulated sinusoidally at a range of frequencies using a piezoelectric objective actuator (P-725.4CD, Physik Instrumente) by up to 10*μm* at different frequencies (1Hz, 0.5 *Hz*, 0.2*Hz*, 0.1*Hz*, see Methods for details).

**Figure 4.**
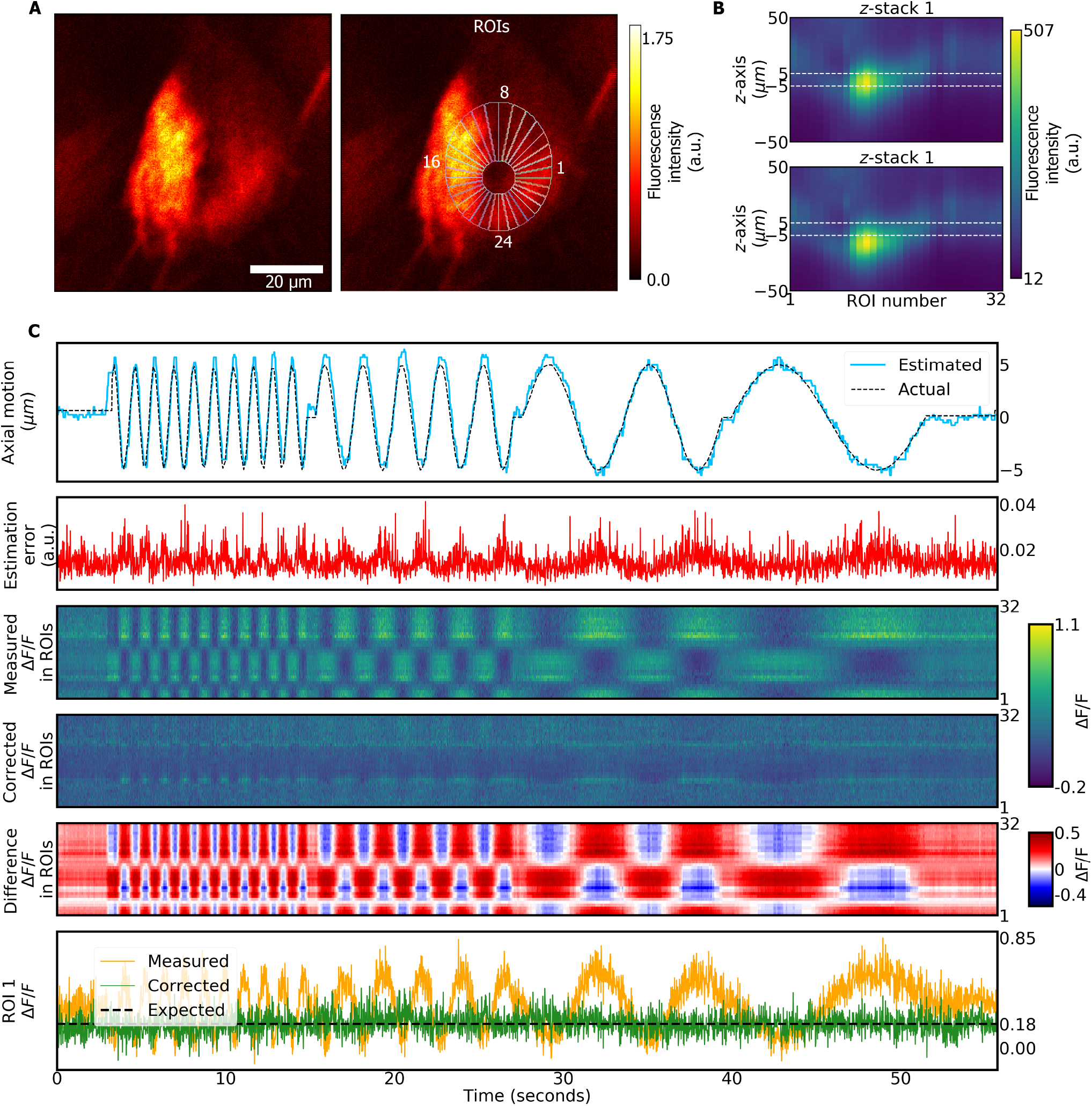
Axial motion estimation and correction in GFP labeled neurons. **A** Left side: average over all frames of the experiment. Right side: 32 ROIs are defined. **B** Stacks recorded from each ROI with first (top) and second (bottom) beam, respectively. **C** Top row: actual and estimated axial motion. Second row: cost function error of motion estimation (see Methods for details). Third row: measured fluorescence changes in each ROI. Fourth row: corrected fluorescence changes in each ROI. Fifth row: difference between corrected and measured fluorescence changes for each ROI. Bottom row: measured, corrected, and expected changes in fluorescence for ROI 1 (representative for all ROIs).

**Figure 5.**
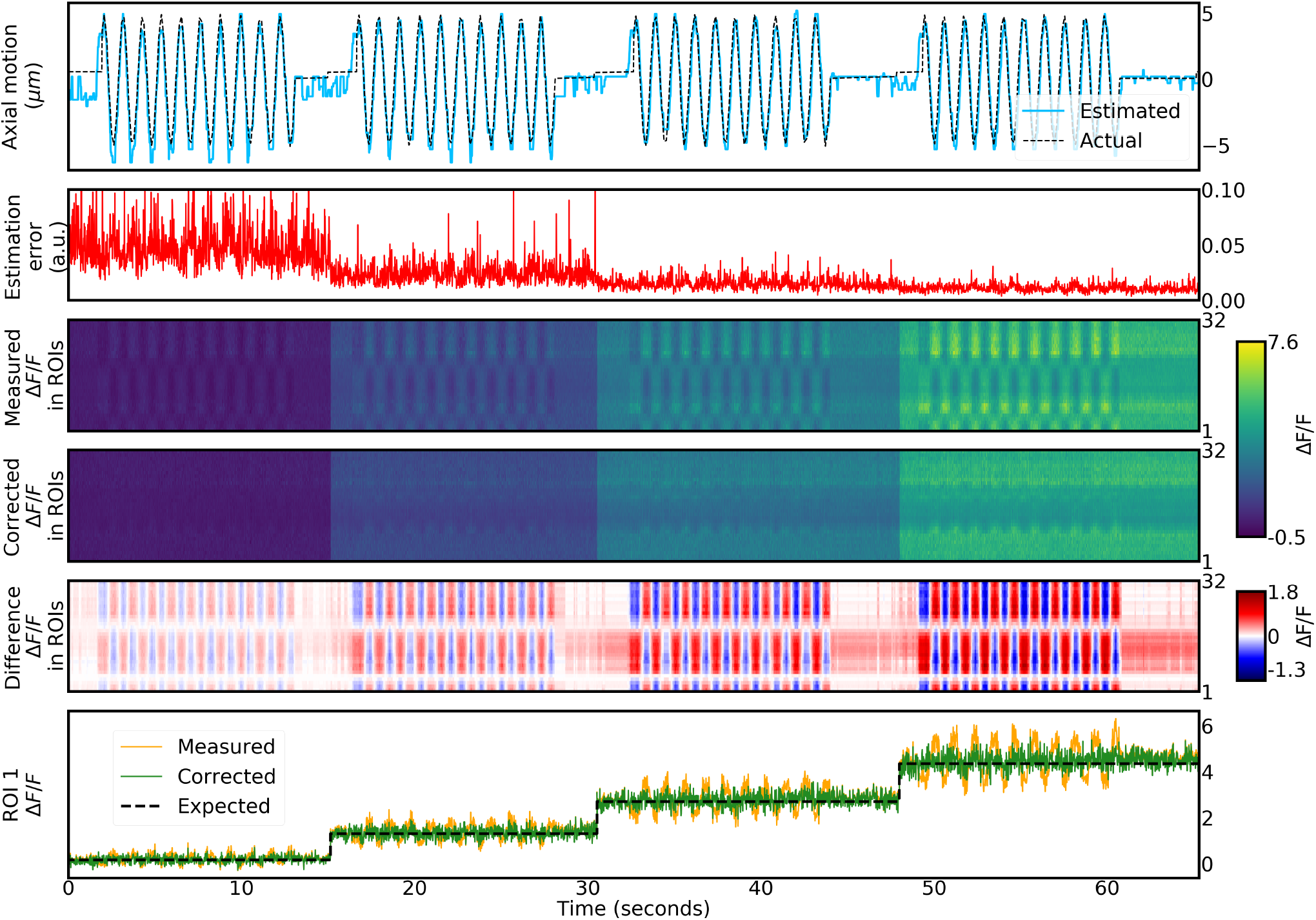
Axial motion estimation and correction in GFP labeled neurons at different laser powers, simulating changes in neural activity. The definition of ROIs and the recorded stacks are the same as in Fig. 4A and B, respectively. Top row: comparison between actual and estimated axial motion. Second row: cost function error of motion correction algorithm. Third row: measured changes in fluorescence in each ROI. Fourth row: corrected changes in fluorescence in each ROI. Fifth row: difference between corrected and measured changes in fluorescence for each ROI. Bottom row: comparison between measured, corrected and expected changes in fluorescence for ROI 1 (see Methods for details).

The motion correction algorithm was applied for estimating axial motion from data simultaneously recorded in two focal planes as follows. At the beginning of the experiment a total of 10 *z*-stacks were recorded with both beams. Each z-stack contained a total of 400 slices covering 100*μm* along the *z*-axis in steps of 0.25*μm*. The resolution of each slice was 256 × 256 pixels and images were recorded at 60Hz. Even though there was no lateral brain motion during these experiments, the piezoelectric objective actuator produced lateral displacements at different z-positions. Therefore, we defined a template, consisting of the 10 first images acquired during the experiment, and all images, including those of the *z*-stacks, were aligned with respect to this template using a phase correlation algorithm (as implemented in Skimage^27^, see Methods). After alignment, the 10 *z*-stacks were averaged and a total of *N* = 32 ROIs were defined along the dendritic toroidal structure as described previously^22^ (Fig. 4A, right side).

The intensities recorded in the *z*-stacks were further filtered using a median filter with a window size of 20 slices to remove noise. The filtered intensities for all ROIs are shown in Fig 4B for both beams.

Axial motion in the time series data was estimated by computing the value of the log-likelihood function (Eq. (35)) in all 400 slices of the *z*-stacks with a time window of size *σ_f_* = 3. Then, the slice 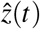 that maximized the log-likelihood function was determined. The resolution of the axial motion estimation was given by the stack’s resolution along the *z*-axis, i.e., 0.25*μm*. As seen in Fig. 4C, top row, the algorithm accurately estimates motion. The cost function error (Fig. 4C, second row, calculated using Eq. (38)) indicates that the accuracy of the estimate is similar across all positions.

The fluorescence intensities were corrected using Eq. (18), and the corrected Δ*F/F* was computed using Eq. (41) (Fig. 4C, fourth row). This was compared to the measured Δ*F/F*, obtained from Eq. (47) (Fig. 4C, third row). The strong changes in measured fluorescence intensity, or motion artifacts, across ROIs in Fig. 4C, third row, are removed after correction and all ROIs show constant fluorescence profiles, as expected for GFP labeled neurons (Fig. 4C, fourth row). This is also seen in the average intensities for a representative ROI (Fig. 4C, bottom row). The offset (from 0) in the corrected Δ*F/F* value is due to an arbitrary definition of the baseline as the fraction of 10 percent lowest intensities (see Supplementary Fig. S2 for details).

### Estimation and correction of controlled sample motion and intensity variations

To estimate at the same time motion related as well as activity induced intensity changes, we varied the focal plane (as described above) together with the laser power, resulting in different levels of fluorescence intensity in a GFP labeled sample (Fig. 5). Four epochs of sinusoidal focal plane modulations at a frequency of 1Hz were combined with increasing the laser power in steps, simulating different neural activity levels. Recordings of time series and *z*-stacks, image preprocessing as well as motion estimation and correction were performed as described above.

Motion was again estimated accurately (Fig. 5, top row). Additionally the algorithm correctly estimated the different motion corrected intensity levels (Fig. 5, third to fifth row). This is also seen in the average changes across all regions of interest (Fig. 5, bottom row). As expected, the cost function error (Fig. 5, second row) decreased with increased signal-to-noise ratio resulting from the increased fluorescence signal. Motion estimation always occurs on a single frame basis (smoothed with a Gaussian kernel with a standard deviation of 3 frames, see Methods), so that faster variations in intensity would result in similar estimates.

### Estimation and correction of controlled motion for *in vivo* two-photon calcium imaging

We next corrected controlled motion of the focal plane in a sample expressing the calcium indicator jGCaMP8f^28^ in the head direction system of the fly^26^. The fly walked in a virtual reality setup on an air supported ball^22^ and the proboscis of the fly was fixed to prevent uncontrolled sample motion. Controlled focal plane motion was again generated by actuating the objective by up to 10*μm* at different frequencies (1Hz, 0.5Hz, 0.2Hz, 0.1Hz). Images (z-stack and time series) were recorded, corrected for lateral motion, and filtered as described above and motion correction proceeded along the same steps (Eqs. (35), (18), (41), and (47)). Again, *N* = 32 ROIs were defined (Fig. 6A, right side) and Fig. 6B shows the intensity profiles of all ROIs for both beams.

**Figure 6.**
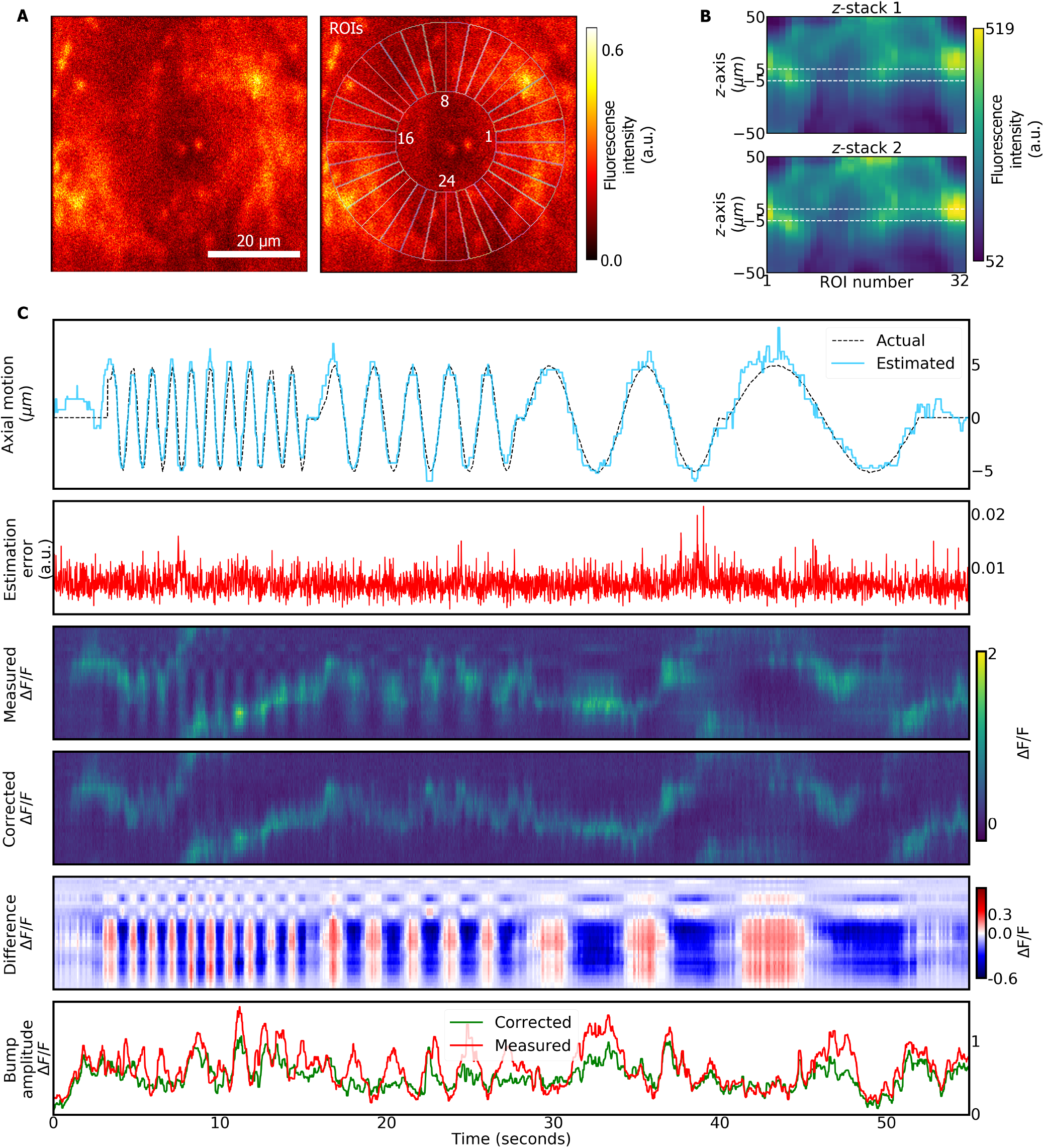
Motion estimation and correction in wedge neurons labeled with jGCaMP8f during controlled axial motion. **A** Left side: average over all frames. Right side: definition of 32 ROIs. **B** *z*-stacks recorded by the first beam (top) and second beam (bottom). **C** First to fifth row: same as Fig. 4C. Bottom row: measured and corrected fluorescence signals (bump amplitude, see Methods for details).

The estimated axial motion is compared with the actual motion of the piezoelectric objective actuator in Fig. 6C, first row. The measured Δ*F/F* as well as the corrected Δ*F/F* values are shown in Fig. 6C, third and fourth rows. The difference between the corrected and measured Δ*F/F* is shown in Fig. 6C, fifth row. The estimation error (Eq. (38)) indicates that motion was tracked with similar quality across all axial positions. The last row in Figs. 6C compares corrected and measured bump amplitudes (see Methods), which was smoothed using a Gaussian filter with a standard deviation in the kernel of 5 frames. Motion correction was able to remove motion artifacts, resulting in a smooth movement of the bump across the ellipsoid body as expected^26^.

### Estimation and correction of brain motion for *in vivo* two-photon calcium imaging in behaving animals

Finally, motion was corrected in a naturally moving brain during in vivo calcium imaging in a fly walking in a virtual reality setup^7, 22^, without fixing the fly’s proboscis or removing muscles in the head. Motion estimation and correction proceeded as described in the previous section.

We measured brain motion of up to 10*μm* (Fig. 7C, top row) and corrections induced changes in the values of Δ*F/F* of up to 40%, as seen in the difference (Fig. 7C, sixth row) between uncorrected and corrected Δ*F/F* (7C, fourth and fifth row, respectively). The bump amplitudes of the corrected and measured Δ*F/F* are shown in Fig. 7, bottom row, and were computed using Eq. (2) and smoothed using a Gaussian filter with standard deviation of 5 frames.

**Figure 7.**
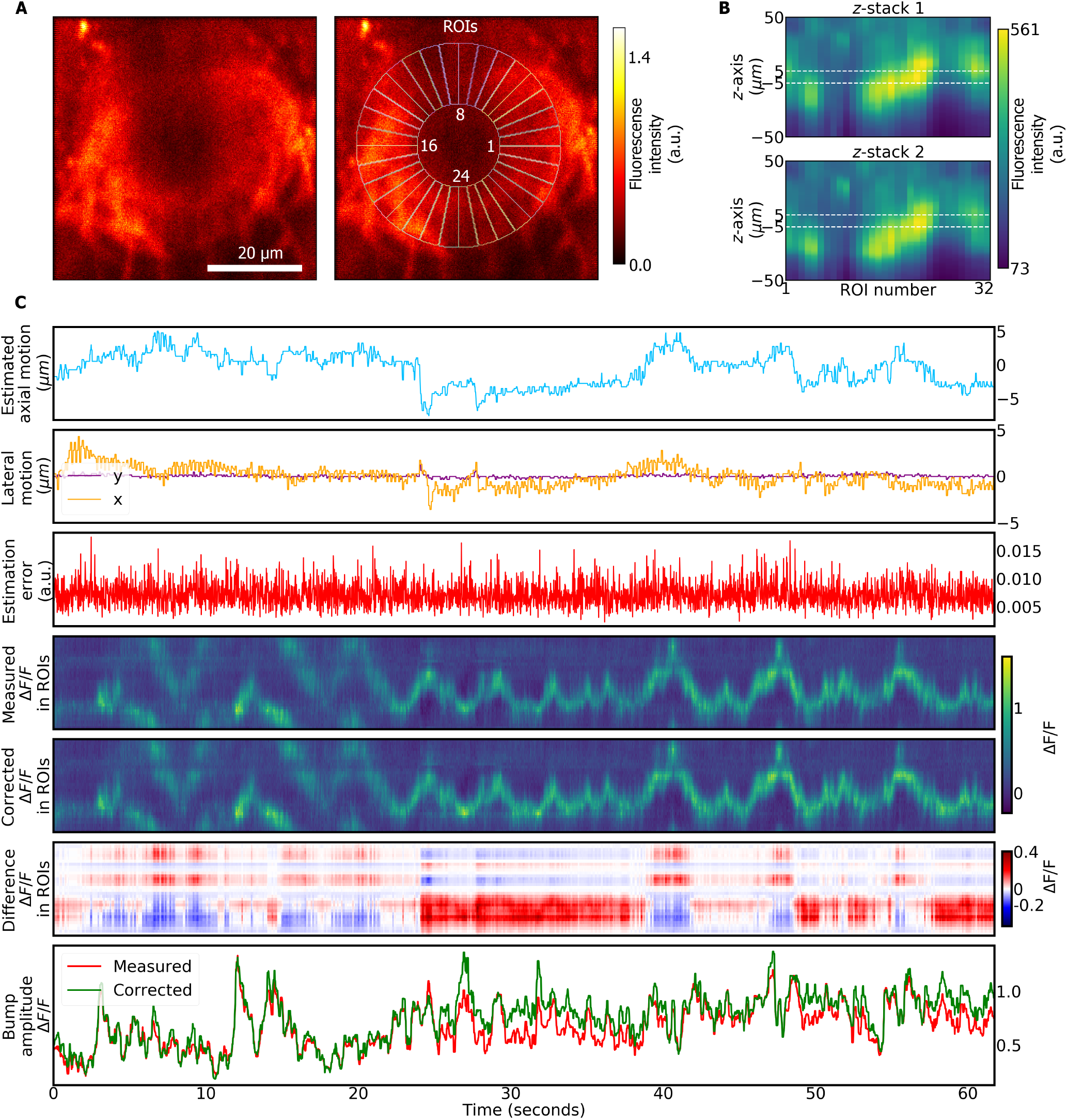
Motion estimation and correction in wedge neurons labeled with jGCaMP8f. **A** Left side: average over all frames. Right side: definition of 32 ROIs. **B** Stacks recorded by the first beam (top) and second beam (bottom). **C** First to sixth row: same as Fig. 4C, additionally including lateral motion estimated from computationally aligning frames (second row). Bottom row: measured and corrected fluorescence signals (bump amplitude, see Methods for details).

### Algorithm for imaging with four simultaneously recorded focal planes

The developed technique can be extended to more than two focal planes (for example four beams as in^18^). Fig. S3 shows a simulation with four axially separated beams. A single ROI was simulated for simplicity (Fig. S3A, top) and the resulting *z*-stacks for all four beams are shown in Supplementary Fig. S3A, bottom. Using more than two beams allows for example increasing the axial resolution or axial range.

## Discussion

We developed a method for correcting motion artifacts for two-photon calcium imaging that result from axial sample movement. The method relies on simultaneous multi-plane imaging with at least two axially offset focal planes^17^. As verified with simulations (Figs. 2, 3 and S3), by controlled motion of the focal plane (Fig. 4, 5, 6), as well as by controlled changes in fluorescence intensity (Fig. 5), the developed optimization algorithm can estimate as well as correct motion with non-ratiometric indicators in isolated structures in the imaging volume.

The method was applied for correcting motion artifacts that are observed when imaging *in vivo* in the brain of fruit flies. Since motion can be corrected without additional mechanical stabilization (Fig. 7), such as removing muscles or immobilizing the proboscis^7^, the approach simplifies imaging in flies and improves the viability of the preparation. Since brain motion in the fly as observed here (about 10*μm*, see Fig. 7) lies in a similar range as observed in other species^4^, the technique could also similarly be applied in other animals.

Compared with motion correction that relies on recording time series of *z*-stacks^12,13^, the approach developed here benefits from high frame rates (for example 60 Hz at 256 × 256 pixels in each plane). Compared with closed-loop correction^4,14–16^, the microscope does not require custom modifications and is compatible with multi-beam scanning as implemented in commercially available scanning software (Scanimage^13^). Further, the developed method does not use an independent tracking fiducial, such as a fluorescent bead or cell body.

That motion is estimated directly from the sample labeled with calcium indicator, leads however to the following two conditions: First, changes in fluorescence due to calcium activity need to occur with a constant gain factor axially along the entire ROI (see Supplementary Fig. S1 for illustration). Given that controlled motion is accurately corrected in a sample expressing jGCaMP8f (Fig. 6), this is a valid approximation for the sample used here. Second: the structure of interest needs to be isolated in the imaging volume at all times. This means that no (partial) structures can move in and out of the imaging volume. Since the algorithm relies on measuring intensity changes along the optical axis, such appearing or disappearing features would lead to erroneous corrections.

Further, the algorithm requires that each ROI must be visible at all times in both focal planes. This condition can be satisfied by adjusting the separation and axial extension of the beams such that the moving object is always within their axial range. Here, beam conditions were optimized for imaging with commonly used GAL4 fly lines labeling subsets of neurons in the central brain. We used two axially extended beams, since we imaged from a relatively large structure. However, the technique could equally be applied for smaller structure with diffraction limited beams. Beam separation and parameters could additionally be adjusted on a sample by sample basis using for example electrically tunable lenses or spatial light modulators. The method can also be similarly extended to more simultaneously recorded focal planes as shown in simulations (Fig. S3) which could be used to extend the axial range or resolution. Additionally, the algorithm requires that lateral motion is corrected first using standard motion correction algorithms and for reliable image alignment therefore sufficient intensity is needed in each channel. Similarly, the ROIs must be defined over a sufficient number of pixels so that the intensity distribution in each ROI is approximately normal.

Overall, the developed method allows correction of lateral and axial brain motion in two-photon calcium imaging experiments from isolated structures in the imaging volume, such as sparse expression patterns in the brain of *Drosophila,* with non-ratiometric indicators.

## Acknowledgements

We would like to thank Giacomo Bassetto for helpful discussions and comments on the algorithm and Ivan Vishniakou for comments on the manuscript.

## Methods

### Microscope setup

The setup was described in^22^. Briefly, two axially offset Gaussian beams were used to record in two different focal planes using temporal multiplexing^17^. Temporal multiplexing was performed using Scanimage in photon counting mode^13^. The diameter and collimation of each beam was adjusted with two lenses (Thorlabs achromatic doublets), while the two beams were linearly and orthogonality polarized and delayed by 2*m*, as shown in Fig. 1B. The beams were focused at the samples with a power of 6*mW* for each beam by underfilling the objective of the microscope (16X Nikon CFILWD Plan Fluorite Objective, 0.80 NA, 3.0 mm WD). This created elongated beam profiles along the axial direction (Fig. 1). The full width at half maximum was adjusted to 9*μm* and 16*μm* (computed from the standard deviations: FWHM = 2.4σ), respectively, and the peaks of the two beams were separated by 7*μm*.

### *Drosophila* preparation

Imaging in behaving flies we performed as described in^22^. Briefly, We used 7-10 days old female flies, either expressing GFP or GCaMP8f in wedge neurons (R60D05-GAL4). We performed laser surgery to remove the cuticle over the brain. The cuticle and underlying tissue were then removed either with a microrobotic arm or manually under a dissection scope^22^. Finally, a drop of glue was placed on the opening (DETAX, Freeform, 02204). The flies expressing GCaMP8f were left to recover over night^22^, while flies expressing GFP were imaged directly after surgery.

For the fly expressing GFP, as well as for the fly in Fig. 6 the proboscis was fixed with wax to prevent brain motion^7^. For imaging, flies were glued to a cover slide ((22 mm × 22 mm, thickness No. 1, Cat. No. 631-0124, see^22^ for details of the preparation).

### Extracting head direction from calcium signals

We computed the trajectory of the bump of activity along the ROIs, *b*(*t*), based on the ROI position with maximum fluorescence value of the corrected Δ*F/F*:

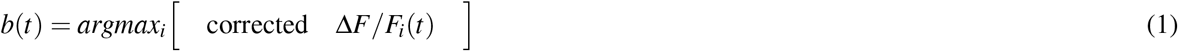

We then used this trajectory to compute the bump amplitude in both, measured and corrected Δ*F/F*, in the ROIs along the bump trajectory:

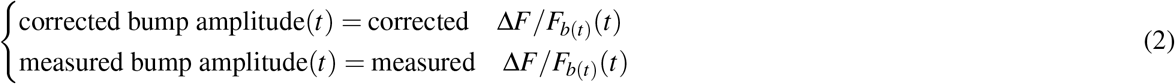

### Motion estimation and correction algorithm

The motion estimation and correction algorithm is shown schematically in Fig. 1A and proceeded along the following steps:

- **1)** First, we recorded a total of 10 *z*-stacks of the sample with each beam, using the Fast *z* mode in Scanimage with a piezoelectric objective actuator. The *z*-stacks were obtained over 100*μm* along the *z*-axis in steps of 0.25μ, at a resolution of 256 x 256 pixels at 60H*z* frame rate. Stacks were recorded simultaneously with the two beams. The time to record the 10 stacks was approximately 67 seconds.
- **2)** We recorded time series of the sample simultaneously in two focal planes at 60H*z*, with the same resolution and zoom as the stacks, providing a total of *T* frames for each beam. Lateral motion correction (i.e., along the *x* and *y*-axis) was performed using the phase correlation algorithm from Skimage^27^ in Python. We defined a template consisting of the first 10 frames of the two-plane time series recordings and computed the offset of each subsequent frame in the time series with respect to this template^9,22^. Additionally, all frames in the recorded stacks were similarly aligned with respect to their respective templates and the aligned 10 stacks for each beam were finally averaged.
- **3)** We defined *N* = 32 regions of interest (ROIs) corresponding to the structural arrangement of wedge neurons^22^. The intensity of each ROI *i* was computed for each beam, *I*_1,*i*_(*t*) and *I*_2,*i*_(*t*), at every recorded frame, *t* = 1,..., *T*, by summing over all pixels within the ROI. We also computed the intensity of each ROI *i* in each averaged *z*-stack, producing a total of 400 slices of each ROI *i* profile recorded by each beam along the *z*-axis, 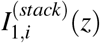 and 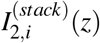.
- **4)** For each frame *t*, we estimated axial motion of the sample, 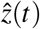, by obtaining the *z* position in the stacks that maximized Eq. (35). An estimation error, *E*(*t*) was additionally computed using Eq. (38).
- **5)** The motion estimation is used to remove axial motion-related changes in the measured intensities *I*_1,*i*_(*t*) and *I*_2,*i*_(*t*), using Eq. (39). Finally, the corrected change in fluorescence in each ROI *i*, Δ*F/F_i_*, is computed according to Eq. (41).

The previous steps are also summarized in Algorithm 1.

#### Algorithm 1: Motion estimation and correction algorithm

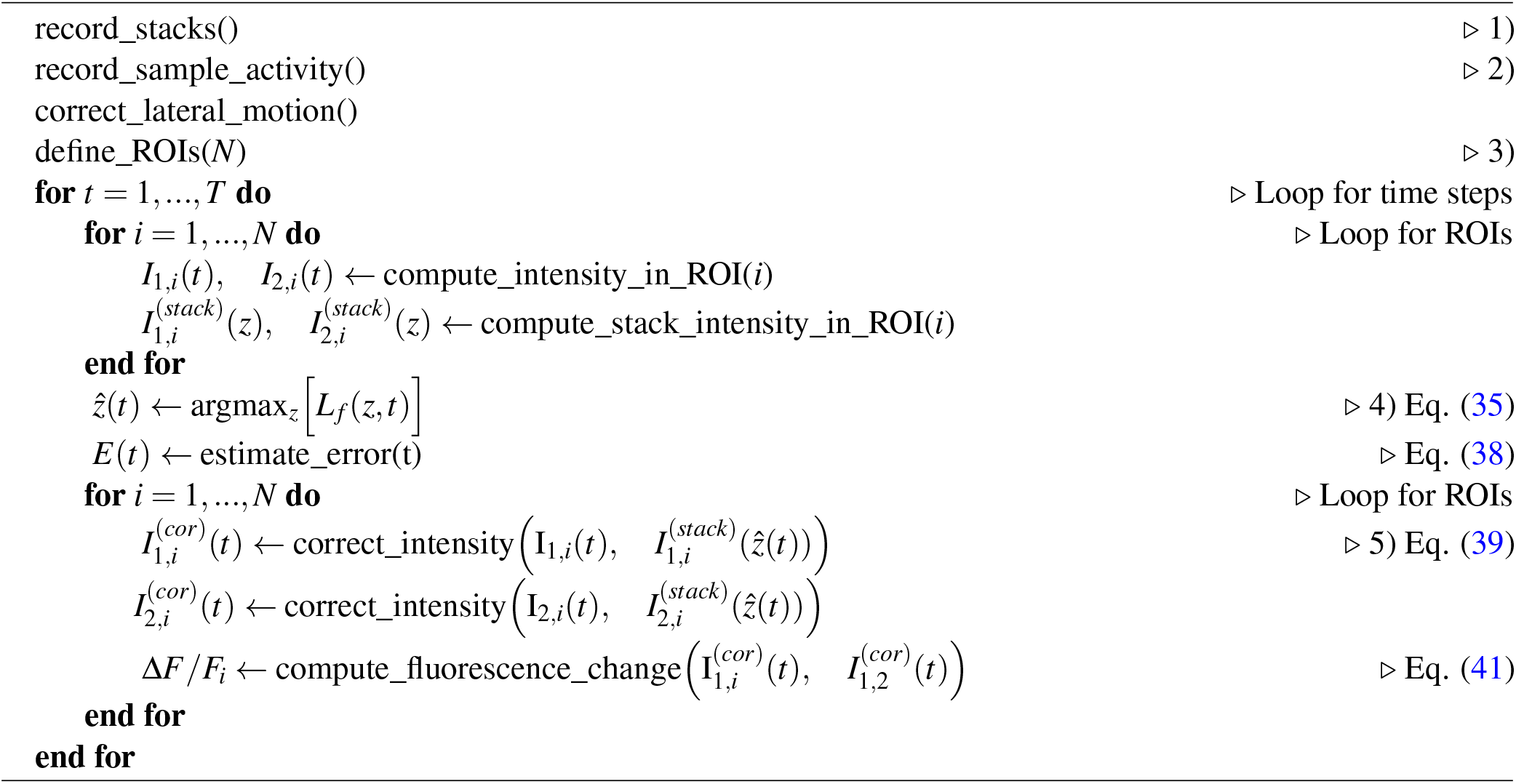

### Analytical approximation of motion correction algorithm for a single ROI

While motion correction was performed numerically as described in Algorithm 1, we here first explain the operation of the algorithm using an analytical approximation, assuming continuous time and *z*-axis, first for a single ROI defined on a 3D sample in the absence of noise.

A single ROI is a voxel extended along the *z*-axis, which is described as a function *e*(*A,z*) representing the number of fluorescent proteins, for example GFP or GCaMP, inside the ROI at a given *z* position. The function *e*(*A, z*) changes over time depending on a variable *A, A* = *A*(*t*), representing neural activity in each ROI. Generally, the ROI function, *e*(*A, z*), could have any form. As an example, Fig. S1A shows a ROI function defined by the following Eq.:

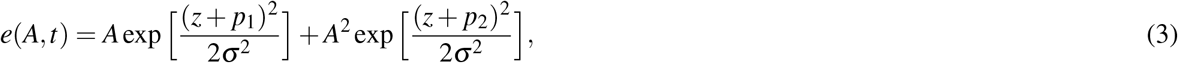

where *p*_1_ = −5*μm* and *p*_2_ = 5*μm* represent the center of two Gaussians with standard deviation *σ* = 2.5*μm*. In this case the activity *A* changes the Gaussian peaks by different amplitudes (Fig. S1A), meaning that fluorescence changes heterogeneously at different *z* positions.This could for example be the case if two different dendrites of a neuron receive inputs with different strengths.

To simplify the problem of motion estimation and correction, we can approximate the ROI function to first order with a Taylor series around *A* = 0:

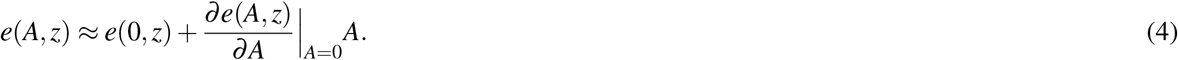

The derivative of *e*(*A, z*) with respect to *A* and evaluated at *A* = 0 is a function that only depends on *z*, which we call *ρ* (*z*). On the other hand, *e*(0, *z*) is the baseline fluorescence indicator distribution along the *z*-axis which is not modulated by the activity *A* of the ROI. We assume for simplicity that *e*(0, *z*) is negligible, assuming that there is an activity baseline, A^0^, much larger than *e*(0,*z*):

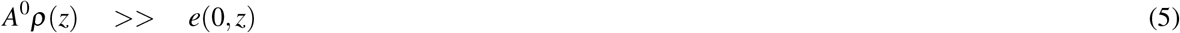

With this approximation, we can write the ROI function as:

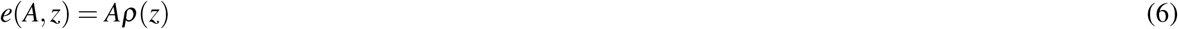

The function *ρ* (*z*) describes a density of fluorescence proteins along the *z*-axis, while *A* increases activity inside the ROI homogeneously. In the following we refer to *ρ* (*z*) as the ROI density while *A* is called the ROI activity. An example of Eq. (6) is given in Fig. S1B, where *ρ (z)* is defined by the following expression:

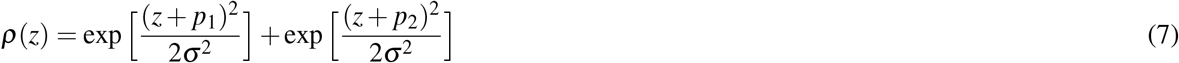

In addition, we assume that the ROI undergoes axial (along the *z*-axis) motion, described with a time-dependent variable, Δ*z*(*t*), which changes the offset of the ROI density along the *z*-axis. Assuming no distortion in the sample during axial motion, the ROI density remains constant but moves along the *z*-axis. Therefore, the ROI function is finally approximated as:

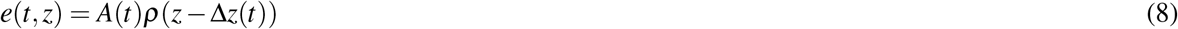

Eq. (8) allows us to separate the ROI function, *e*(*t, z*), into two parts, one which depends on the activity of the ROI, *A*(*t*), which is independent of the *z*-axis, and another one that depends on the motion of the sample, *ρ* (*z* – Δ*z*(*t*)).

This first order approximation, which was valid in the biological samples used for the experiments, allows us to develop an algorithm to estimate and correct the axial motion of the sample The goal of the algorithm is to estimate axial motion, Δ*z*(*t*), as well as to extract the activity, *A*(*t*).

The ROI activity is measured using two beams described by two functions, *g*_1_ (*z*) and *g*_2_(*z*), which represent the shape of the beams along the *z*-axis. For the algorithm to work, these two beams must have different profiles along the *z*-axis (to provide unique information about the sample) and must be integrable, meaning that their integral must be finite. In practice, we assume that these are Gaussian beams centered at different positions along the *z*-axis, while their width along the *z*-axis and power can be different. At each time, *t*, both beams excite the sample with a time delay (on the order of the fluorescence lifetime), and produce two independently measured fluorescence signals.

The intensity of each fluorescence signal associated with each beam, *I*_1_(*t*) and *I*_2_(*t*), is the integral of the excitation of each beam within the ROI along the *z*-axis:

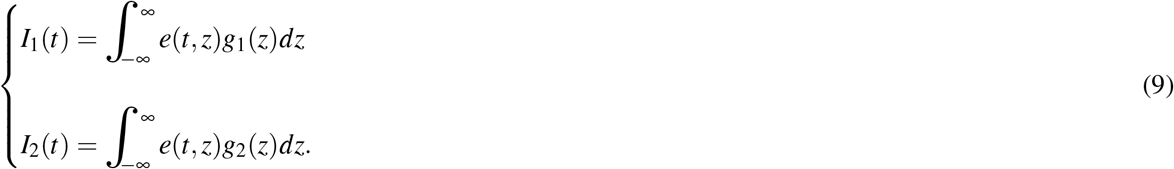

Since the ROI activity, *A*(*t*), is independent of *z* (Eq. (8)), we can write the measured intensities as:

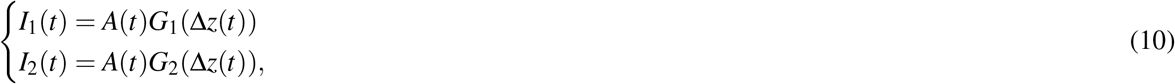

where *G*_1_ and *G*_2_ are two functions defined by the integral over the excitation of each beam at the ROI density, *ρ*(*z* – Δ*z*(*t*)), and only depend on the axial motion of the sample, Δ*z*(*t*). Eq. (10) describes how changes in the measured intensities, *I*_1_(*t*) and *I*_2_(*t*), can have two different contributions: the activity of the ROI and the axial motion of the sample.

For estimating axial motion of the sample, Δ*z*(*t*), a calibration step is first performed. At the beginning of the experiment (*t* = 0), a *z*-stack is recorded with each beam by moving the two beams simultaneously and continuously along the *z*-axis. We assume that the ROI activity and axial motion of the sample, *A*(0) and Δ*z*(0), do not change when recroding the stack. Moreover, and without loss of generality, we define the origin of the axial motion while recording the stacks so that Δ*z*(0) = 0. In practice, the assumption that *A*(0) and Δ*z*(0) do not change is achieved by averaging over several stacks (see Results with biological samples expressing GFP or GCaAMP8f). The algorithm however works with arbitrary activity *A*(0) during *z*-stack acquisition. Since moving the beams is equivalent to moving the sample in the opposite direction while leaving the beams fixed, the stacks are defined as follows:

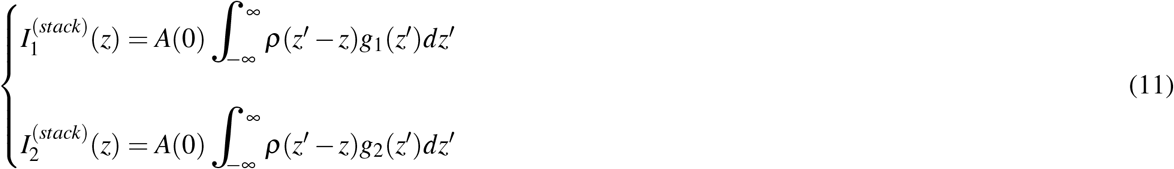

This equation can be expressed as

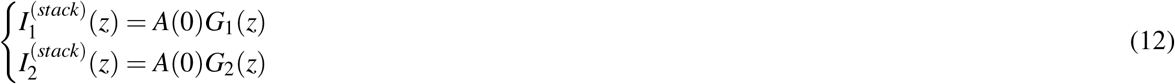

The ratios *I*_1_ (*t*)/*I*_2_(*t*) and 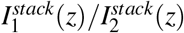 do not depend on the activity and we use this to define the following cost function,

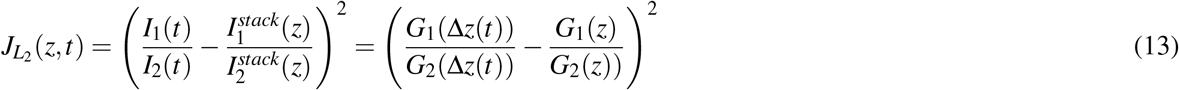

The cost function *J*_*L*1_ (*z*) is minimized at time *t* when *z* = Δ*z*(*t*). This optimization is used to find the axial motion of the sample, Δ*z*(*t*), using the stacks. We will show in the following sections (Eq. (34)) that the shape of this cost function arises from a expectation–maximization approach. To take into account that the axial motion of the sample is correlated in time, we modified Eq. (13), with a Gaussian filter over time:

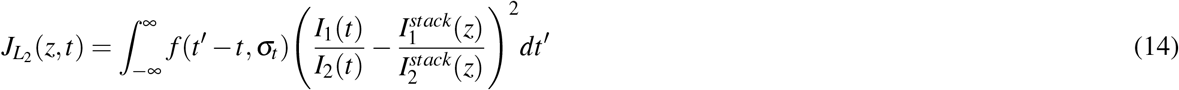

where *f* (*t* – *t*’, *σ*) is a Gaussian kernel centered at time *t* with standard deviation *σ_t_*, defined as:

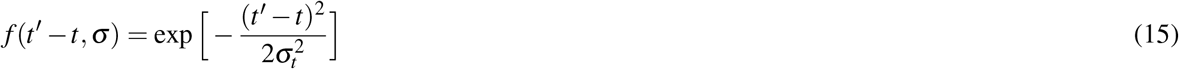

Eq. (14) provides a filtered estimate of the axial motion of the sample by weighting contributions from several intensity measurements. This cost function assumes that changes in axial motion within the Gaussian kernel are small. The standard deviation, *σ_t_*, is a manually-set parameter that provides the size of the time window considered for the axial motion estimation.

To find the axial motion of the sample at any time *t*, we find the *z*-position in the stack, 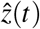, that minimizes the cost function.

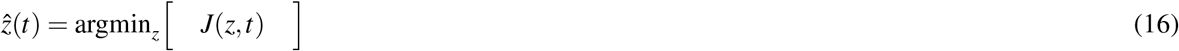

The value of the cost function evaluated at the estimated displacement, 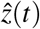, provides a measure for the quality of the optimisation result at time *t*.

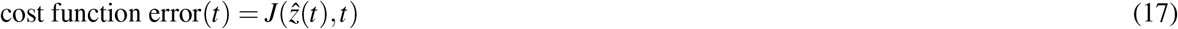

Once the the motion of the sample is estimated, we correct the measured intensities by dividing the intensity recorded by each beam by its corresponding stack, evaluated at the estimated slice 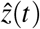 at any time *t*:

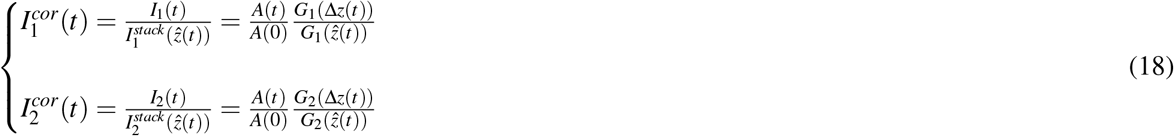

This correction eliminates the contribution of the axial motion of the sample from the measured intensities if the axial motion estimation is correct, i.e. 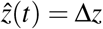, leaving only the contributions made by changes in the activity *A*(*t*). Finally we can compute the relative change in fluorescence, Δ*F/F*, using the corrected intensities:

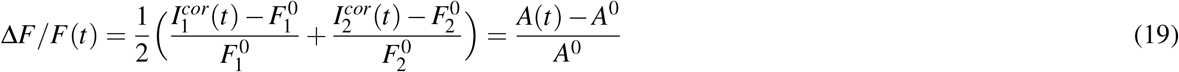

where 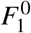 and 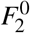 are the baselines of the intensities 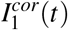 and 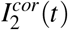 respectively, corresponding to the ROI activity baseline, *A*^0^. The baselines 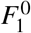 and 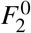 are obtained by computing the mean value of the lowest 10% values of the corrected intensities, 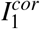 and 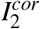, respectively.

### Simulation of a moving single ROI

To demonstrate how this algorithm works for a single ROI, we simulated a single ROI along the *z*-axis described by Eq. (6) with an activity density defined as

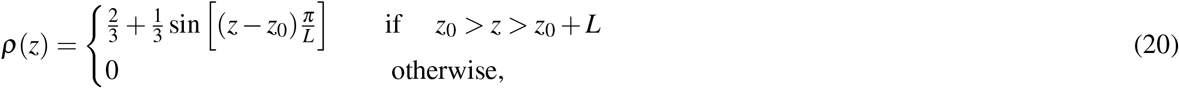

where *z*_0_ = – 5*μm* and *L* = 10*μm*. The ROI undergoes axial motion over time, as defined by Eq. (8) and shown in Fig. 2B, described by the following Eq.:

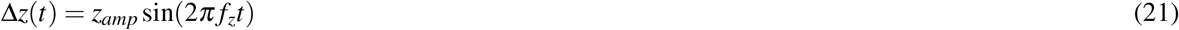

where *z_amp_* = 5*μm* and *f_z_* = 5H*z* are the amplitude and frequency of a simulated sinusoidal axial motion. The activity of the sample, *A*(*t*), is modeled by a differential Eq.:

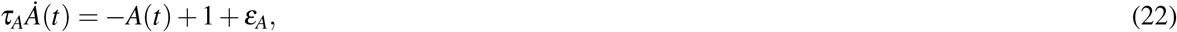

where *τ_A_* = 10*ms* is a time constant, and *ε_e_* is a stochastic input to the sample that produces spikes along the simulation with 0.5% probability and positive amplitude.

To record the activity of the sample, we defined two beams with Gaussians axial profiles, according to the following equation:

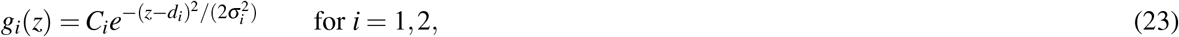

where *C*_1_ = 1.25*a.u*. and *C*_2_ = 1.5*a.u*. define the maximum beam power and *σ*_1_ = 2*μm* and *σ*_2_ = 3*μm* are the widths (standard deviations) along the *z*-axis. The two beams are offset in axial direction, *d*_1_ = –2.5*μm* and *d*_2_ = 2.5*μm*. The shape of the beams is shown at the top of Fig. 2A, together with the ROI function at time *t* = 0, *e*(0, *z*).

A *z*-stack of the sample at time *t* = 0 is defined in the absence of axial motion, Δ*z*(0) = 0, and activity is defined as *A*(0) = 1. The stack is simulated by recording the activity of the sample while moving the beams along the the *z*-axis from –25*μm* to 25*μm* in steps of 0.05*μm*, resulting in a convolution between the beams and the sample according to Eq. (11) (Fig. 2A). Only a single stack without averaging is used for simulations.

Next, combined sample motion and activity are simulated for 1000 milliseconds. The equation for the activity, *A*(*t*), is solved using forward Euler with a time step of *dt* = 0.001 milliseconds. The intensity of each beam is given by Eq. (9). The result of the simulation is shown in Fig. 2C, where the first row shows the moving sample and its activity, and the second row shows the intensities recorded in each beam.

To estimate sample motion, 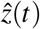, at each instant *t*, we computed the value of the cost function (Eq. (14)) at all *z* positions of the stacks (from –25μ to 25μ in steps of 0.05μ). Finally, motion at time *t*, Δ*z*(*t*), is estimated according to Eq. (16), by finding the position 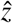 that minimizes the cost function across all *z* values. The third row in Fig. 2B shows the real versus the estimated motion of the sample. The error of the optimisation process is calculated using Eq. (17) and shown in the fourth row of Fig. 2.

Finally, the recorded intensities are corrected using the estimated axial motion and Eq. (18), and changes in fluorescence intensity, Δ*F/F*, are computed from the corrected intensities, according to Eq. (19). The last row in Fig. 2 shows the actual and the corrected changes in fluorescence, respectively. The actual Δ*F/F* is calculated according to the following Eq.:

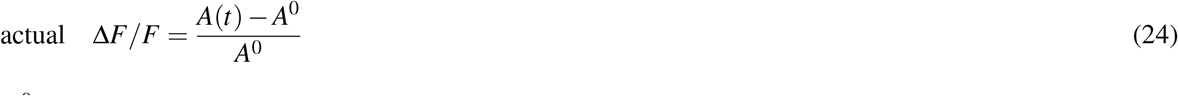

where *A*^0^ is an activity baseline calculated as the average of the lowest 10% values of *A*(*t*).

### Algorithm for motion estimation and correction for several ROIs with noise

We now extend the approach developed above to a sample with multiple ROIs. In addition we now consider that the measurements are noisy. We assume rigid three-dimensional sample motion, with negligible elastic deformations or changes in orientation.

We record a total of *T* pair of images, *M*_1_ (*t*) and *M*_2_(*t*), one for each beam, for *t* = 1,..., *T* and assume that all images are aligned, that is, corrected for lateral motion. We define *N* ROIs on the images and each ROI *i* describes an area on the images containing ni pixels. The sum of the values of all the pixels of each image inside this area, 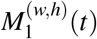 and 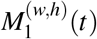, where (*w, h*) ∈ ROI *i*, provides a pair of intensity measurements of the ROI *i* in each pair of images at time *t*:

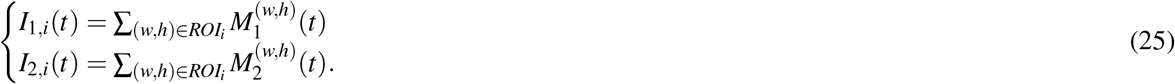

The fluorescence signals detected by the PMT are subjected to shot noise and the intensities in each pixel follow independent Poisson distributions. If the number *n_i_* of pixels in each ROI *i* is large, each pair of intensity measurements approaches the following normal distributions:

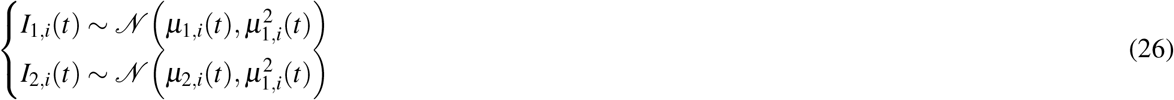

where μ_1,*i*_ and μ_2,*i*_ are the means of the normal distribution for each beam, respectively.

Using the first-order Taylor expansion (Eq. (4)), we can approximate each ROI function *i* by the following expression:

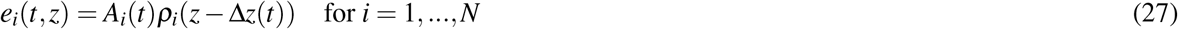

where *A_i_*(*t*) and *ρ_i_*(*z* – Δ*z*(*t*)) are the activity and density of each ROI *i*, respectively, during the acquisition *t*. We again assume that the ROI moves in the axial direction, given by Δ*z*(*t*). Then, we can approximate the means of the intensity distributions, μ_1,*i*_(*t*) and μ_2,*i*_(*t*), as

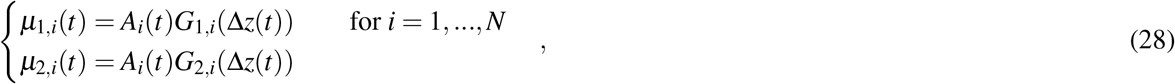

where *G*_1,*i*_(Δ*z*(*t*)) and *G*_2,*i*_(Δ*z*(*t*)) are functions defined by the integral of the excitation of each beam at each ROI density, *ρ_i_* (*z*)_*i*_.

We now take the ratio of the intensities for each ROI *i* at each acquisition *t*, *I*_1,*i*_(*t*)/*I*_2,*i*_ (*t*), since the ratio of the mean values μ_1,*i*_(*t*)/μ_2,*i*_(*t*) does not depend on the ROI acivity, *A_i_*(*t*). The ratio of these two normal variables is well approximated by a normal distribution around the mean of the ratio, μ_1,*i*_(*t*)/μ_2,*i*_(*t*), if the means are positive and the coefficients of variation, 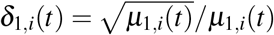 and 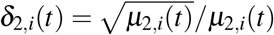 are smaller than 0.1^29^.

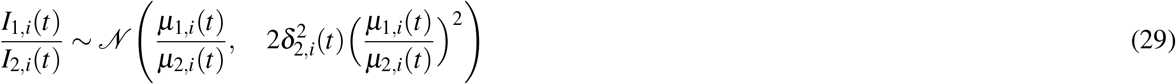

Assuming that all ROIs undergo the same axial motion, Δ*z*, we can compute the log-likelihood for the ROIs with respect to their axial position, Δ*z*, at acquisition *t*:

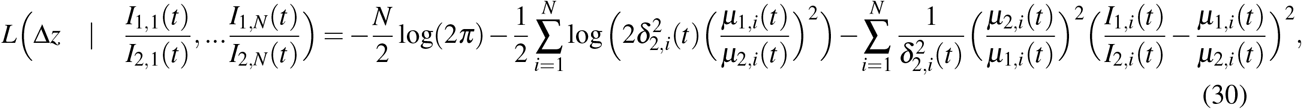

where the mean of the ratio distributions depends only on the axial motion Δ*z*(*t*), and not on the activity of the ROIs:

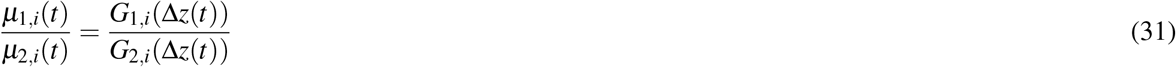

To compute the ratio of the means for each ROI *i*, we used *z*-stacks recorded at the beginning of the experiment, *t* = 0. Each slice in each stack is aligned to remove lateral motion using phase correction^27^, and the intensity for each ROI *i* in each *z*-slice is computed, 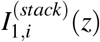 and 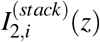. During the acquisition of the stack, both axial motion and activity of each ROI are considered constant, while the noise in the intensities of the slices in the stacks are assumed zero. Therefore, the intensity of each ROI in each slice of the stacks is given by the following equation:

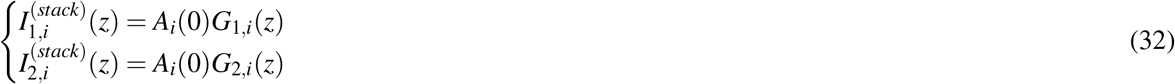

In practice, the previous assumptions are achieved by averaging several stacks and applying a median filter (see Results). We can now approximate the mean of the ratio distribution by the ratio of the intensities in the stacks at the slice Δ*z*:

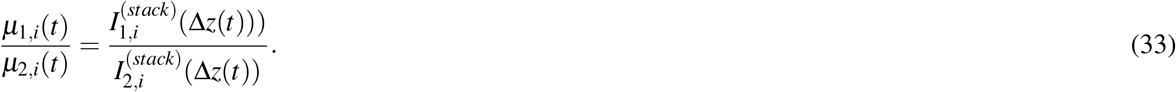

Finally, we can write the log-likelihood (Eq. 30) using the stacks as a function of *z* at each acquisition, *t*:

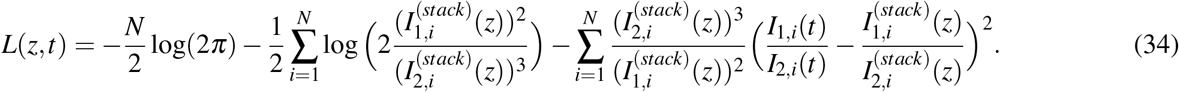

Note that *L*(*z, t*) has a quadratic difference between the ratio of the stacks and the ratio of intensities (right side of Eq. (34)), similar to the cost function defined for a single ROI (Eq. (13)), which was minimized due to the change in sign. The log-likelihood function, however, weights (left factor on the third term in Eq. (34)) and offsets (second term in Eq. (34)) this quadratic difference, taking into account more strongly ROIs that are less noisy to estimate the axial motion, i.e., the intensity of those ROIs with lower values (and therefore lower standard deviation in their distributions).

Since axial motion is correlated in time, we use a Gaussian filter similar to Eq. (14)) in the log-likelihood:

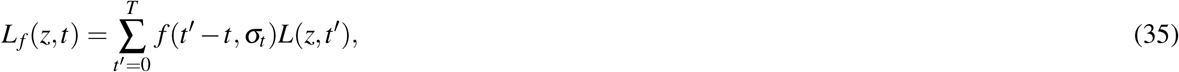

where the kernel *f* (*t*’ – *t, σ_t_*) is defined in Eq. 15. Again, the variable *σ_t_* is manually set, and defines the size of the time window considered for the sum of log-likelihoods.

Since the *z*-stacks have a finite number of slices, we can compute the value of *L_f_* (*z, t*) numerically for each slice *z* and then estimate the axial motion of the sample, 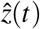, at each time *t*, from the slice *z* that maximizes the log-likelihood function, *L_f_* (*z, t*):

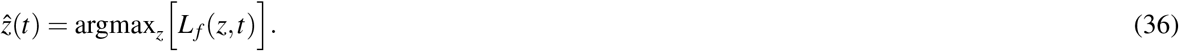

We provide an estimation error, *E*(*t*), by computing the standard deviation of the difference between the ratio of recorded intensities and the ratio of the means:

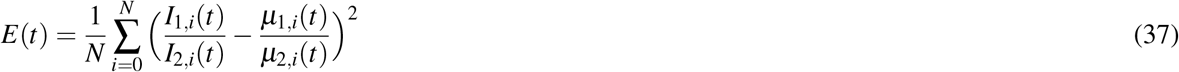

and using Eq. (33), this error can be expressed as:

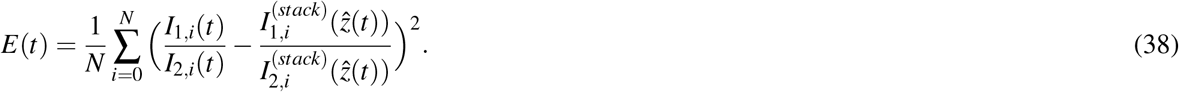

Next, we correct the measured intensities from each beam using the estimated axial motion and extending Eq. (18) to *N* ROIs:

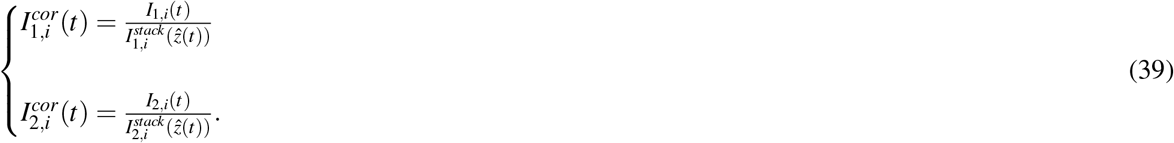

This correction eliminates the axial motion contribution from the intensity measurements. However, note that this correction inherits the noise from the measured intensities *I*_1,*i*_(*t*) and *I*_2,*i*_(*t*). According to Eq. (26), both intensity corrections are obtained according to the following normal distribution:

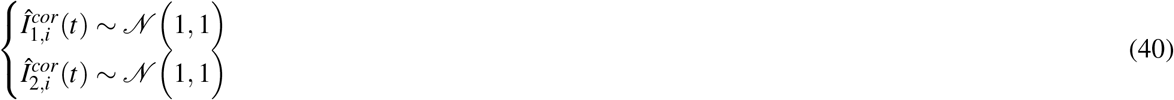

Finally, we compute the change in fluorescence for each ROI *i* as follows:

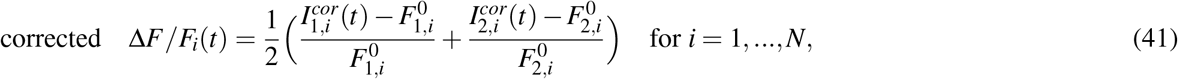

where 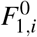 and 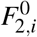 are the baselines of the intensities, 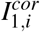 and 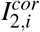 that correspond to the activity baseline of each ROI, 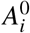. As for the case of a single ROI, we compute the baselines 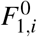 and 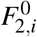 from the average of the 10% lowest values of 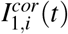 and 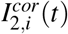 respectively, for each ROI *i*.

### Simulation of several moving ROIs

We demonstrate how the algorithm works for a simulation of *N* = 32 moving ROIs with noise. In this simulation it is assumed that lateral motion of a 3D sample is already corrected and the ROIs are defined. The density of each ROI is defined along the *z*-axis according to the following equation:

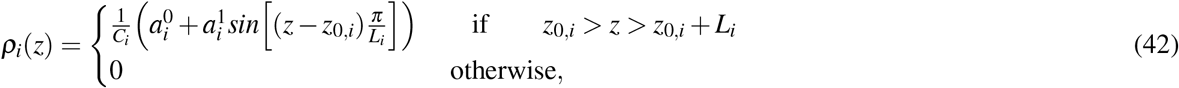

where 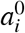 and 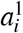 are random coefficients for each ROI obtained from an uniform distribution in the range [1,2] and *z*_0,*i*_ and *L_i_* are a random origin and random length for each ROI, sampled from a uniform distribution in the range of [–2, –5] and [10,15], respectively. The constant *C_i_* normalizes the density of each ROI so that *ρ_i_*(*z*) is in the range of [0,1]. Fig. 3A, top, shows the density of each ROI used in the simulation.

The motion of all ROIs is modeled as in the simulation in section using Eq. (21) with a frequency of *f_z_* = 8*Hz*. The activity of each sample is modeled as:

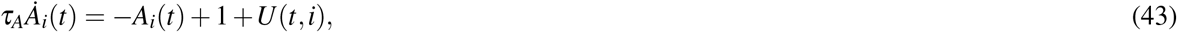

where *τ_A_* = 10*ms* is the time constant and *U*(*t, i*) is an input defined as a rotating Gaussian along the ROIs:

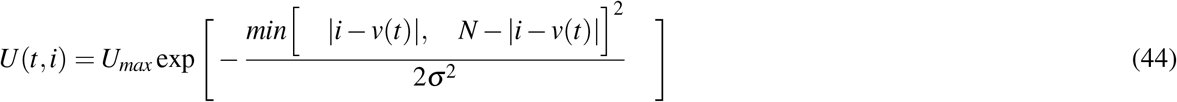

where *U_max_* = 2 is the amplitude, *σ* = 2 is the width of the Gaussian and *v*(*t*) rotates the input with respect to the ROIs at a frequency of *f*_u_ = 10*Hz*, according to

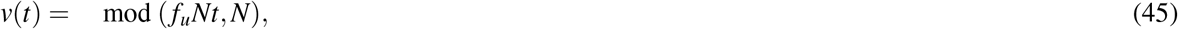

where mod (·, ·) indicates the modulo operation. The activity of the ROIs is again solved using forward Euler with a time step of *dt* = 0.001 millisecond. An example of ROI 1 undergoing axial motion while receiving the rotating input is shown in Fig. 3A, bottom, while the rotating input in all the ROIs during the simulation is shown in Fig. 3C, fourth row.

Two beams, defined again by Eq. (23), were used to record the fluorescence change in all ROIs. At time *t* = 0 we recorded a *z*-stack using 1000 slices (from –25*μm* to 25μ in steps of 0.05*μm*), assuming no axial motion of the sample, Δ*z*(0) = 0. The recorded *z*-stacks are shown in Fig. 3B. Note that the activity during the *z*-stack acquisition is higher around the ROI 16. The algorithm can estimate and correct the axial motion with arbitrary activity *A_i_*(0) in the *z*-stack.

After defining the stacks, the intensities at time *t* in two simultaneous planes for each ROI i are simulated, *I*_1,*i*_(*t*) and *I*_1,*i*_(*t*), with changing axial position and ROI activity. The intensities in each plane are sampled from a Gaussian distribution, given by Eq. (26), where the mean values *μ*_1,*i*_(*t*) and *μ*_2,*i*_(*t*) are computed using Eq. (28).

To estimate axial motion of the sample, Δ*z*(*t*), we computed the value of the log-likelihood function in Eq. (35) at all 1000 *z* positions of the stacks, at any time *t*, using a time window of size *σ_t_* = 3. The estimated axial motion is then obtained by the slice in the *z*-stacks, 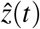, that maximizes the log-likelihood function. The first row of Fig. 3C shows the estimated compared to the actual axial motion during the simulation, while the estimation error, computed by Eq. (38), is shown in the second row of Fig. 3C.

Next we corrected the measured intensities from each beam and each ROI using the estimated axial motion and Eq. (39). The change in fluorescence of the corrected intensities, corrected Δ*F/F_i_*(*t*), is computed using Eq. (41) and shown in Fig. 3C, fourth row. This is compared with the actual Δ*F/F_i_*(*t*) (Fig. 3C, fifth row), which is computed as:

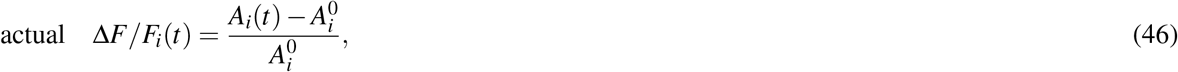

where 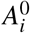 is the baseline activity of each ROI *i*, obtained from the average of the 10% lowest values of *A*(*t*). Fig. 3C, third row, shows the change in fluorescence measured without motion correction. This measured Δ*F/F_i_*(*t*) is obtained by the following Eq.:

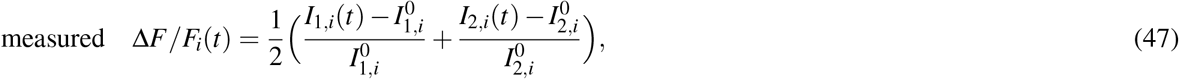

where 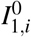 and 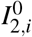 are the baseline measured intensities for each beam, obtained by the mean value of the 10% lowest values of *I*_1, *i*_(*t*) and *I*_2,*i*_(*t*), respectively. These baseline values are affected by axial motion and therefore changes in fluorescence appear much larger (Fig. 3C, third row) when the sample is out of focus due to the resulting low intensities. Further, the corrected Δ*F/F* is larger than the actual Δ*F/F* (Fig. 3, fourth and fifth row). This is due to noise in the measurements, which produces lower baselines values, 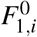 and 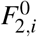. These lower values follow from averaging over the lowest 10% values of the corrected intensities (see for example Fig. S2B) instead of taking the mean value of the distribution.

### Simulation for motion estimation and correction with four simultaneously recorded focal planes

Here, we extend the motion correction algorithm to four beams using simulations. For simplicity we consider only a single ROI. The ROI density is defined again by Eq. (20). Four different Gaussian beams are defined, according to the following equation:

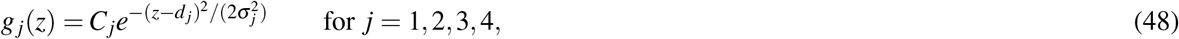

where *C*_1_ = 1, *C*_2_ = 1.5, *C*_3_ = 2 and *C*_4_ = 0.5 are the maximum beam powers, and σ_1_ = 3, σ_2_ = 3, σ_3_ = 2 and σ_4_ = 3 are the beam widths (standard deviations) along the *z*-axis. Each beam is offset from the previous one by 3.33*μm* (*d*_1_ = –5, *d*_2_ = —1.67, *d*_3_ = 1.67 and *d*_4_ = 5). The beam profiles, as well as the ROI densities, are shown in Fig. S3A, first row.

First, a stack is recorded by continuously moving each beam long the *z* axis, from —25*μm* to 25μ in steps of 0.05μ. The stack obtained for each beam is shown in the second row of S3A. We assume that the ROI undergoes axial motion over time, defined by Eq. (21), while its activity changes according to (22). We ran the simulation for a total of 1000 milliseconds using forward Euler with a time step of *dt* = 0.001. Both activity and axial motion of the ROI are shown in the first row of Fig. S3B. At each time step *t* each beam recorded intensity from the sample according to the following equation:

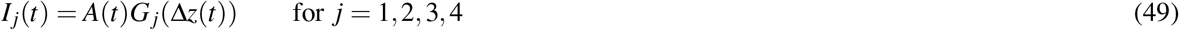

where the functions *G_j_*(Δ*z*(*t*)) represent the integration of the excitation of each beam *j* at the ROI density. The intensities recorded by each beam are shown in the second row of Fig. S3.

To estimate the axial motion of the ROI, we extended the cost function for two beams (Eq. 14) and defined the following cost function for four beams:

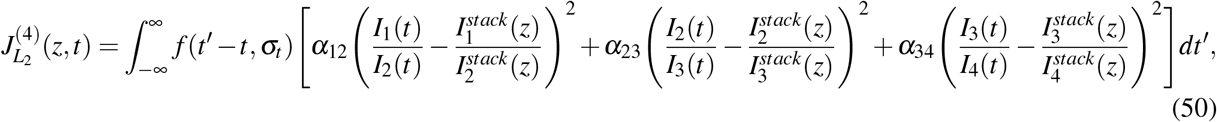

where *f* (*t*’ – *t, σ_t_*) is the Gaussian kernel defined by Eq. (15), and the parameters *α*_12_, *α*_23_ and *α*_34_ are computed at each time step *t* defined as:

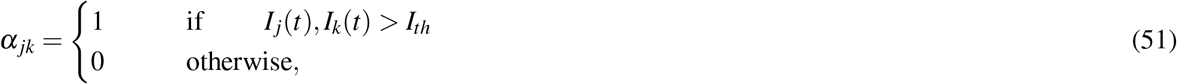

**Figure S1.**
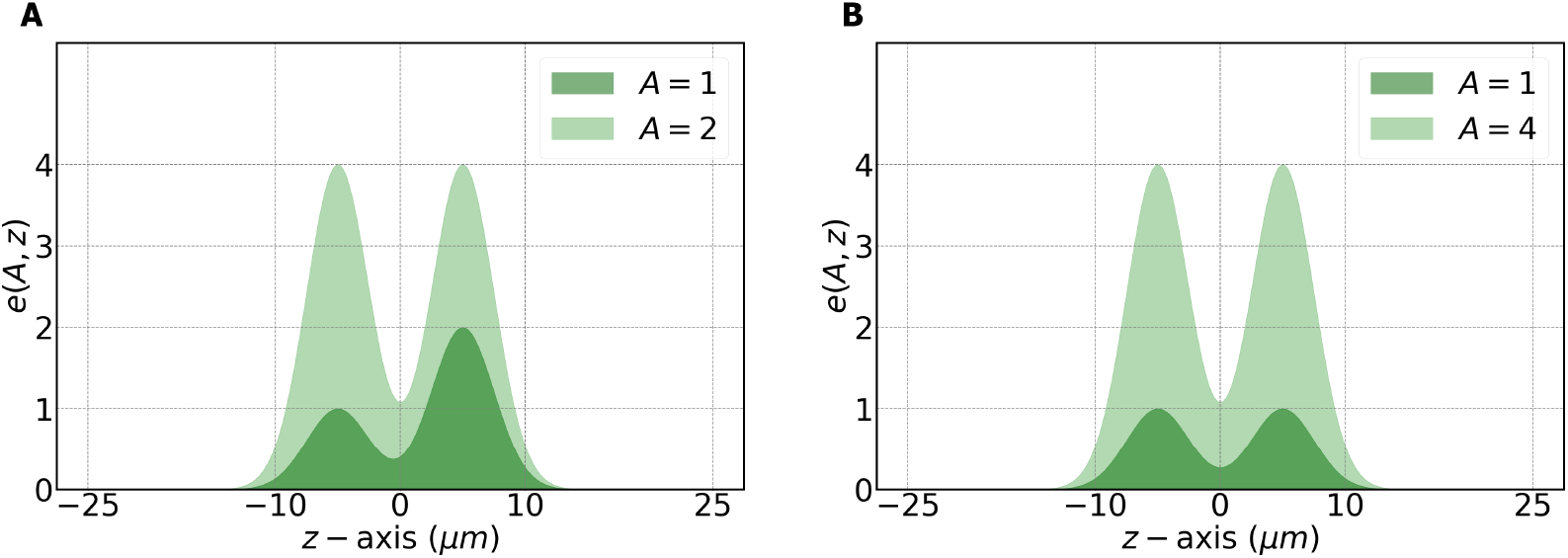
Example of two different ROI functions. *A* ROI function defined by Eq. 3, where the activity *A* modulates heterogeneously the function along the *z*-axis. Such modulation can not be corrected with the developed algorithm. **B** ROI function defined by Eq. 7, where the activity *A* homogeneously modifies the ROI function along the *z*-axis. An underlying assumption of the algorithm is that activity is modulated in such a homogeneous fashion from baseline.

where *I_th_* = 0.1 is a minimum intensity threshold. The parameters *α_jk_* are used to only take into account measured intensities at *t* if the ROI is within the field of view of beams *j* and *k*. When the intensity recorded by one of the beams is lower than *I_th_*, the ROI is considered outside of the z-field of view of the beam and therefore its contribution to the cost function of Eq. (50) is discarded by the *j* parameters.

We estimated the axial motion of the single ROI by finding the slice in the stacks, Δ*z*(*t*), at any time step *t*, that minimized the cost function (50). Both the estimated and actual axial motion are shown in Fig. S3B, third row. The cost function error, shown Fig. S3B, fourth row, is given by the value of the cost function evaluated at the estimated axial motion, 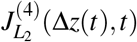.

Finally, the intensity of each beam is corrected using the following expression:

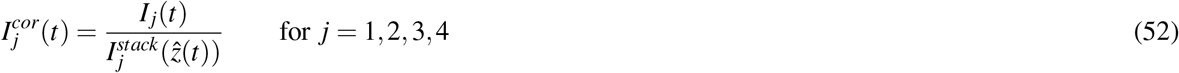

and the corrected fluorescence change, Δ*F/F* is computed from the corrected intensities:

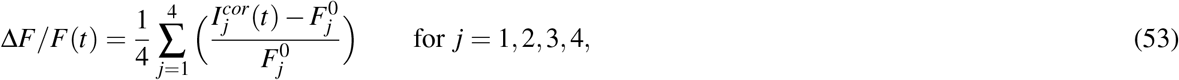

where 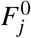 is the baselines of the intensity 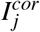, computed from the average of the 10% lowest values of 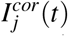 for beam *j* = 1,..., 4. The last row of Fig. S3B shows the corrected, actual, and measured Δ*F/F*. The actual Δ*F/F* is computed according to Eq. 24. The measured Δ*F/F* is the fluorescence change that would be measured assuming no motion correction, given by:

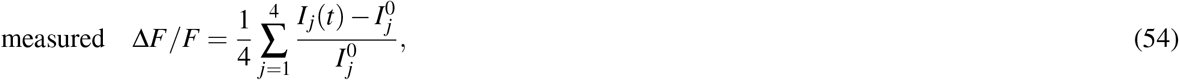

where 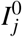 is the baseline measured intensity, obtained by the mean value of the 10% lowest values of *I_j_* (*t*) for each beam *j* = 1,...,4.

**Figure S2.**
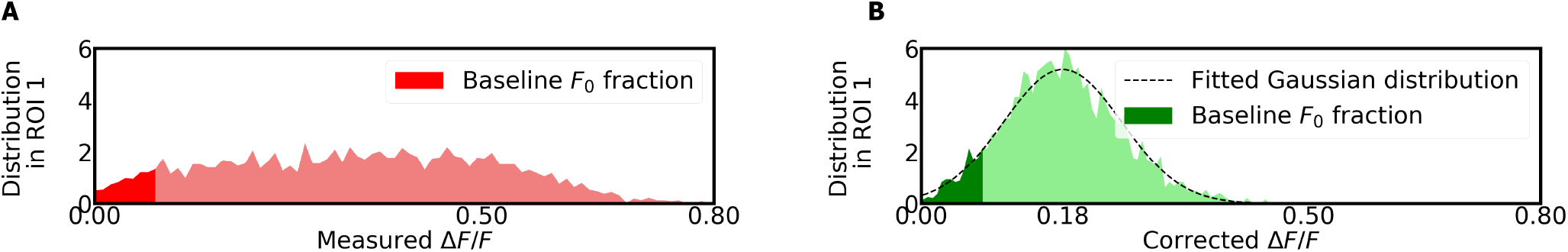
Distributions of fluorescence changes for ROI 1 in the experiment shown in Fig. 4. **A** Distribution of the measured Δ*F/F* for ROI 1. This distribution is affected by axial motion. The distribution of the measured Δ*F/F* for ROI 1 is not Gaussian due to the axial motion **B** Gaussian distribution of the corrected Δ*F/F* for ROI 1. Since the fluorescence baseline is calculated from the average of 10% lowest values, the mean of the distribution of corrected Δ*F/F* is not centered around 0, as one would expect for GFP (no fluorescence changes).

**Figure S3.**
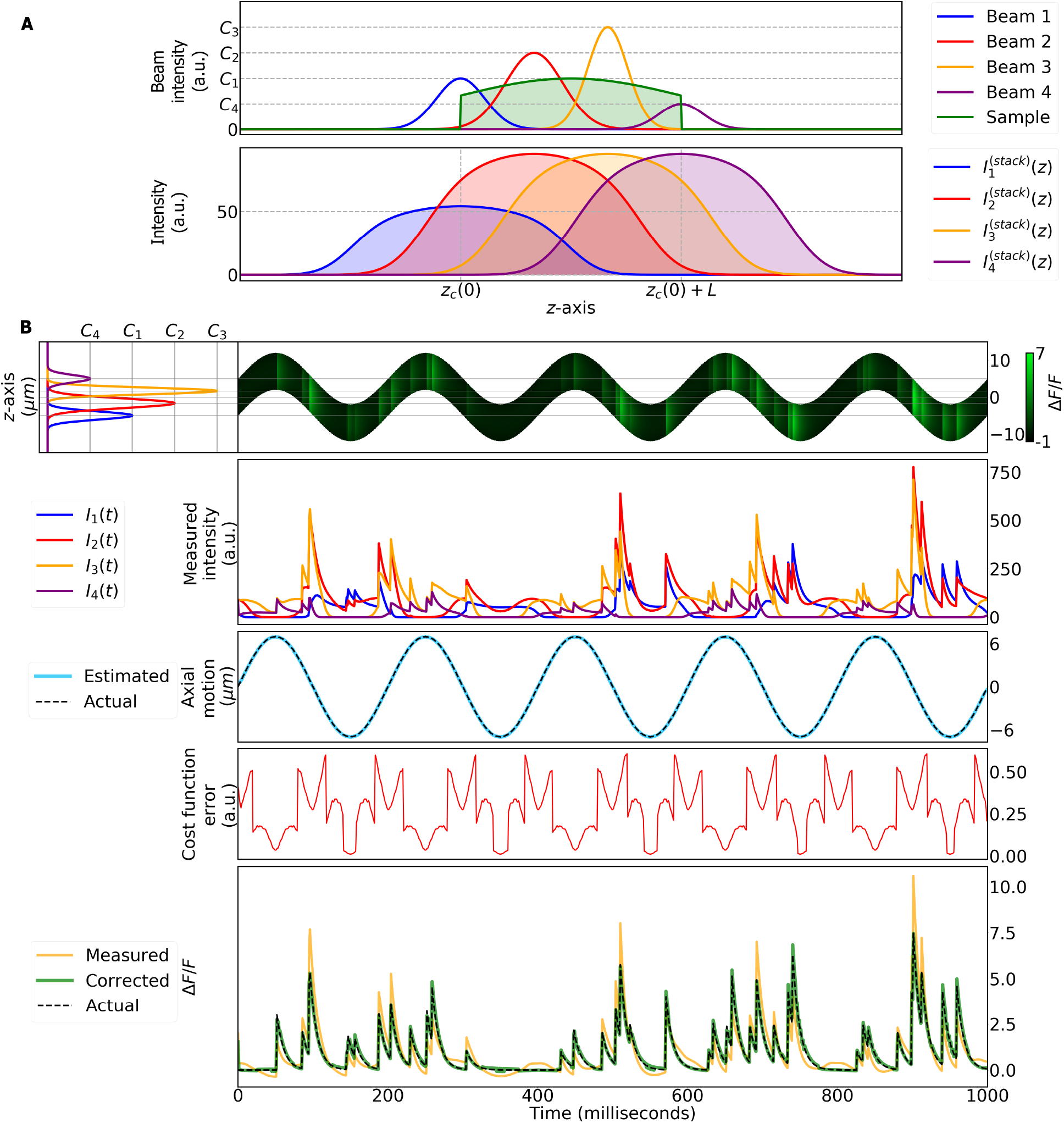
Simulation of a single moving voxel for a configuration with four temporally mulitplexed beams. **A** Top row, ROI density at time *t* = 0 as well as the profiles of the four beams along the *z*-axis. Bottom row, stack of the sample at time *t* = 0 obtained for each beam. **B** First row, left side: profile of the four beams along the *z*-axis. Right side: activity and axial motion of the ROI over time. Second row: intensity measured with each beam over time. Third row: estimated and actual axial motion of the ROI over time. Fourth row: cost function error evaluated at the estimated axial position. Bottom row: measured, corrected and actual activity of the ROI over time.

